# Structural insights into heterohexameric assembly of epilepsy-related ligand–receptor complex LGI1–ADAM22

**DOI:** 10.1101/2025.01.06.631603

**Authors:** Takayuki Yamaguchi, Kei Okatsu, Masato Kubota, Ayuka Mitsumori, Atsushi Yamagata, Yuko Fukata, Masaki Fukata, Mikihiro Shibata, Shuya Fukai

## Abstract

Leucine-rich glioma-inactivated 1 protein (LGI1) is a secreted neuronal protein consisting of the N-terminal leucine-rich repeat (LRR) and C-terminal epitempin repeat (EPTP) domains. LGI1 is linked to epilepsy, a neurological disorder that can be caused by genetic mutations of genes regulating neuronal excitability (*e.g*., voltage- or ligand-gated ion channels). ADAM22 is a membrane receptor that binds to LGI1 extracellularly and interacts with AMPA-type glutamate receptors via PSD-95 intracellularly to maintain normal synaptic signal transmission. Structural analysis of the LGI1–ADAM22 complex is important for understanding the molecular mechanism of epileptogenesis and developing new therapies against epilepsy. We previously reported the crystal structure of a 2:2 complex consisting of two molecules of LGI1 and two molecules of the ADAM22 ectodomain (ECD), which is suggested to bridge neurons across the synaptic cleft. On the other hand, multiangle light scattering, small-angle X-ray scattering, and cryo-EM analyses have suggested the existence of a 3:3 complex consisting of three molecules of LGI1 and three molecules of ADAM22. In the previous cryo-EM analysis, many observed particles were in a dissociated state, making it difficult to determine the three-dimensional (3D) structure of the 3:3 complex. In this study, we stabilized the 3:3 LGI1–ADAM22_ECD_ complex using chemical crosslinking and determined the cryo-EM structures of the LGI1_LRR_–LGI1_EPTP_– ADAM22_ECD_ and 3:3 LGI1–ADAM22_ECD_ complexes at 2.78 Å and 3.79 Å resolutions, respectively. Furthermore, high-speed atomic force microscopy (HS-AFM) visualized the structural features and flexibility of the 3:3 LGI1–ADAM22_ECD_ complex in solution. We discuss new insights into the interaction modes of the LGI1–ADAM22 higher-order complex and the structural properties of the 3:3 LGI1–ADAM22 complex.

**Significance:** The neuronal secretory protein Leucine-rich glioma-inactivated 1 (LGI1) and its receptor protein ADAM22 play a critical role in maintaining normal synaptic signal transmission. Genetic mutations in LGI1 are linked to epilepsy. Structural analysis of the LGI1–ADAM22 complex is crucial for understanding the molecular mechanisms of epileptogenesis and for developing targeted treatments. In this study, we determined the cryo-EM structure of the heterohexameric LGI1–ADAM22 complex and visualized the dynamics of this complex by high-speed atomic force microscopy.

## Introduction

Epilepsy is a prevalent neurological disorder, affecting approximately 1% of the population. Epilepsy is characterized by recurrent, unprovoked seizures, resulting from an imbalance between excitation and inhibition within neural circuits. Mutations associated with epilepsy frequently occur in genes that regulate neuronal excitability through ion channels such as voltage-gated ion channels (*e.g*., K^+^, Na^+^, and Ca^2+^ channels) and ligand-gated ion channels (*e.g*., nicotinic acetylcholine and GABA_A_ receptors) (1-3). Additionally, some epilepsy-related mutations have been identified in genes encoding non-ion channel proteins such as *LGI1* (4-7).

LGI1 is a 60-kDa secreted neuronal protein that consists of the N-terminal leucine-rich repeat (LRR) domain and the C-terminal epitempin-repeat (EPTP) domain (7) (Fig. 1a). Mutations in the LGI1 gene, resulting in incorrect folding and posttranslational modifications, cause autosomal dominant epilepsy with auditorial features (ADEAF) (4-6). For example, the E383A mutant of LGI1 is eliminated by the endoplasmic reticulum and becomes deficient in secretion (8). Meanwhile, the S473L mutation of LGI1 causes epileptiform seizures due to reduced binding to ADAM22, which is a member of the a disintegrin and metalloproteinase (ADAM) family (9, 10) and acts as a receptor for LGI1 in neurons without protease activity (11, 12). ADAM22 is a single-pass transmembrane protein. The ectodomain (ECD) of ADAM22 consists of a metalloprotease-like domain, a disintegrin domain, a cysteine-rich domain, and an EGF-like domain (13) (Fig. 1a). The metalloprotease-like domain interacts with the EPTP domain of LGI1 in the extracellular space (11, 14). In the cytoplasm, the PDZ-binding motif (PBM)-containing C-terminal tail of ADAM22 binds to a synaptic scaffolding protein PSD-95, which regulates the cellular dynamics of α-amino-3-hydroxy-5-methyl-4-isoxazolepropionic acid (AMPA) receptors through binding to Stargazin, an auxiliary subunit of AMPA receptors, in the postsynapse (15, 16). Furthermore, LGI1 forms a complex with the voltage-gated potassium channel (VGKC) through ADAM22/23 (9, 17, 18). As such, the LGI1-ADAM22 complex maintains normal nerve signal transduction through the interaction network including synaptic ion channels (10, 11, 19). Structural analysis of the LGI1-ADAM22 complex is important for understanding the molecular basis of epileptogenesis and for developing therapeutic strategies based on this understanding.

**Figure 1.**
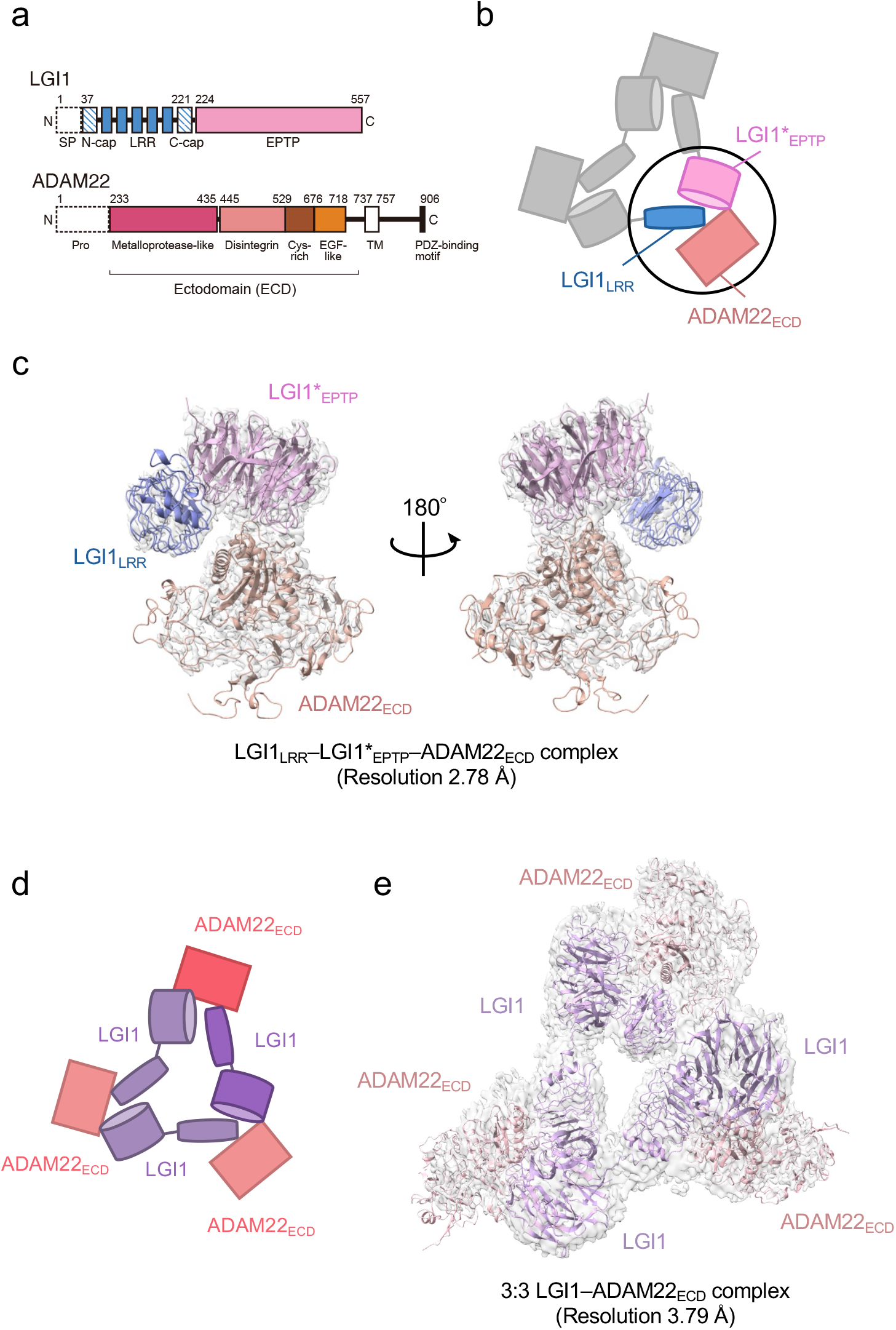
Structure of the LGI1–ADAM22_ECD_ complex. **a** Domain organizations of LGI1 and ADAM22. LGI1 consists of the LRR (blue) and EPTP (pink) domains. The N-terminal secretion signal peptide (SP, enclosed by dotted lines) is removed in the secreted LGI1. The shaded blue boxes represent the N- and C-terminal caps, whereas the filled blue boxes represent the LRRs. The premature form of ADAM22 contains the N-terminal prosequence (enclosed by dotted lines). The mature ADAM22 consists of the metalloprotease-like (magenta), disintegrin (salmon pink), cysteine-rich (brown), EGF-like (orange), transmembrane (TM; white), and cytoplasmic domains. The major ADAM22 isoform has a PDZ-binding motif in the C-terminal of the cytoplasmic domain. **b** Schematic diagram of the LGI1_LRR_– LGI1*_EPTP_–ADAM22_ECD_ complex. The black circle indicates the location of this complex within the 3:3 LGI1-ADAM22 complex. * indicates a distinct molecule. **c** Cryo-EM map and structure of the LGI1_LRR_–LGI1*_EPTP_–ADAM22_ECD_ complex at 2.78 Å resolution. **d** Schematic diagram of the 3:3 LGI1–ADAM22_ECD_ complex. **e** Cryo-EM map and structure of the 3:3 LGI1– ADAM22_ECD_ complex at 3.79 Å resolution.

We previously reported the crystal structure of a 2:2 LGI1–ADAM22_ECD_ complex consisting of two molecules of LGI1 and two molecules of ADAM22_ECD_ (14). The long-axis length of the 2:2 complex is approximately 190 Å, which is comparable to the width of the synaptic cleft. This 2:2 complex structure and the structure-guided study on a mouse model for familial epilepsy suggested that the formation of the 2:2 complex bridges neurons in the synaptic cleft. The results revealed the structural basis of the interaction between the EPTP domain of one LGI1 and the LRR domain of the other LGI1, as well as the interaction between the EPTP domain of LGI1 and the metalloproteinase-like domain of ADAM22 (14). On the other hand, size-exclusion chromatography-multiangle light scattering (SEC-MALS), size-exclusion chromatography-small-angle X-ray scattering (SEC-SAXS), and cryo-electron microscopy (cryo-EM) analyses suggested that three molecules of LGI1 and three molecules of ADAM22_ECD_ bind to each other to form a 3:3 complex at near physiological salt concentrations (14). Similarly to the 2:2 complex, the 3:3 complex might serve as an extracellular scaffold to stabilize Kv1 channels or AMPA receptors in a *trans*-synaptic fashion (9, 17, 19). In addition, the 3:3 assembly in a *cis* fashion on the same membrane might regulate the accumulation of Kv1 channel complexes at the axon initial segment (18, 20). However, no clear evidence to prove these potential mechanistic roles of the 3:3 assembly has been provided, and the three-dimensional structure of the 3:3 complex has not yet been determined.

In this study, the three-dimensional (3D) structure of the 3:3 LGI1–ADAM22_ECD_ complex was determined at a nominal resolution of 3.79 Å by cryo-EM single-particle analysis. We also determined the 3D structure of the LGI1_LRR_–LGI1_EPTP_–ADAM22_ECD_ complex at a nominal resolution of 2.78 Å. These higher-resolution 3D structures provide more detailed interaction mechanisms for the LGI1–ADAM22 higher-order assembly. We also performed high-speed atomic force microscopy (HS-AFM) (21, 22) to directly visualize the molecular dynamics of the LGI1–ADAM22_ECD_ assembly and characterize its structural property. The cryo-EM and HS-AFM results suggest that the 3:3 LGI1–ADAM22_ECD_ assembly does not form a rigid 3-fold symmetric structure but rather a flexible triangular structure accompanying relative motions between each protomer.

## Results

### Cryo-EM single particle analysis of the LGI1–ADAM22_ECD_ complex

As we reported previously, the molar mass of the complex between the full-length LGI1 and ADAM22_ECD_ determined by SEC-MALS at 150 mM NaCl was 356 kDa, corresponding to the 3:3 hexameric assembly of LGI1–ADAM22_ECD_ in solution (14). This is consistent with our initial cryo-EM analysis, where 5% of the reference-free 2D class averaged images showed particles with pseudo-*C*3 symmetry suggestive of the 3:3 assembly. Since the 3:3 complex class was clearly seen and appeared structurally stable, we tried to determine the 3D structure of the 3:3 LGI1-ADAM22_ECD_ complex by single particle analysis but failed due to insufficient numbers of particles of the 3:3 complex. On this basis, we decided to perform a structural analysis of the complex stabilized by chemical crosslinking. We co-expressed His_6_-tagged LGI1 and non-tagged ADAM22_ECD_ in Expi293F cells and purified the complex using Ni-NTA affinity chromatography and gel filtration chromatography at 50 mM NaCl (Fig. S1a,b). Then, fractions containing the LGI1–ADAM22_ECD_ complex were collected and treated with glutaraldehyde to chemically crosslink the 3:3 complex. The sample was purified again by gel filtration chromatography. The chromatogram showed a peak likely corresponding to the 3:3 LGI1–ADAM22_ECD_ complex (Fig S1c,d). One of the peak fractions was collected and subjected to single-particle analysis by cryo-EM. Among 2,766,936 particles picked from 7,625 movies without templates, 1,403,037 particle images were extracted with a box size of 544 pixels (0.752 Å/pixel) and downsampled to a size of 136 pixels (3.008 Å/pixel) by Fourier cropping. The downsampled images were subjected to reference-free 2D classification, which generated images of triangle-shaped particles considered to be the 3:3 LGI1–ADAM22_ECD_ complex (Fig. S2). Other images looked like two particles stacked on top of each other (Fig. S2) or single particles clearly smaller than the 3:3 complex. The stacked particle images might represent the 2:2 complex viewed along the long axis or the 3:3 complex viewed from the side, which were difficult to distinguish. Six classes with 176,443 particles of the putative 3:3 LGI1-ADAM22_ECD_ complex were used as templates for picking particles again. 2,530,790 particle images were extracted, downsampled, and subjected to the second run of reference-free 2D classification, resulting in a set of images similar to that generated in the first run of 2D classification (Fig. S3). Eighty 2D classes with 2,006,398 particles were selected for *ab initio* 3D reconstruction. Six classes of *ab initio* 3D models were generated and then refined by heterogeneous refinement. The resultant 3D map in one class corresponded to a complex comprising LGI1_LRR_, LGI1*_EPTP_, and ADAM22_ECD_ (* indicates a distinct molecule hereafter). Non-uniform refinement using the particle images with the original pixel size yielded a density map at a nominal resolution of 2.78 Å (Figs. 1b,c and S4 and Table S1). This map corresponds to a part of the 3:3 complex, where two-thirds of the complex may adopt various conformations. The 3D map of one other class corresponded to the 3:3 LGI1–ADAM22_ECD_ complex. Non-uniform refinement using the particle images with the original pixel size yielded a density map at a nominal resolution of 3.79 Å (Figs. 1d,e and S4 and Table S1). Using these two maps, we constructed atomic models of the LGI1_LRR_–LGI1*_EPTP_–ADAM22_ECD_ complex and the 3:3 LGI1-ADAM22_ECD_ complex (Table S1). The resolutions of the present cryo-EM analysis (2.78 Å and 3.79 Å) are better than that of the crystal structure of the 2:2 complex (7.125 Å).

### Interaction of the LGI1_EPTP_–ADAM22_ECD_ complex

In the cryo-EM structure of the LGI1_LRR_–LGI1*_EPTP_–ADAM22_ECD_ complex, the LGI1_EPTP_– ADAM22_ECD_ structure is essentially identical to that determined by X-ray crystallography (Fig. 2a). At the interface, Trp398, Tyr408, and Tyr409 of ADAM22 are stacked in a layer and project into the hydrophobic inner rim of the central channel of LGI1_EPTP_, which consists of Leu237, Phe256, Val284, Leu302, Tyr433, Met477, and Phe541 of LGI1 (Fig. 2a). In addition, several hydrogen bonds are formed between LGI1 and ADAM22: Arg330 and Lys353 of LGI1 form hydrogen bonds with Asp405 and Glu359 of ADAM22, respectively, whereas Arg378 of LGI1 forms hydrogen bonds with Ser340 and Thr406 of ADAM22 (Fig. 2b,c and Fig. S5). The hydrogen bond between Asp431 of LGI1 and Lys362 of ADAM22 was also found in the cryo-EM structure (Fig. S5). These hydrogen bonds differ slightly from those in the previous crystal structure: In the previous crystal structure, Lys331 of LGI1 formed a hydrogen bond with the main-chain carbonyl of Asp405 of ADAM22. On the other hand, in the present cryo-EM structure, Lys331 of LGI1 is reoriented and does not form a hydrogen bond (Fig. 2b). Arg378 of LGI1 hydrogen bonds with Ser340 and Glu359 in the crystal structure, while it does with Ser340 and Thr406 in the cryo-EM structure (Fig. 2c). In the crystal structure, Asp431 of LGI1 faces outward and does not form a hydrogen bond with Lys362 of ADAM22 (Fig. S5). These differences in hydrogen bonding may reflect the varying contribution of each interacting residue to the affinity between LGI1_EPTP_ andADAM22_ECD_. As shown in our previous pull-down experiments, mutations of the hydrophobic residues at the interface of ADAM22 (*i.e*., Trp398, Tyr408, and Tyr409) almost or completely abolished binding to LGI1, whereas the E359A or D405A mutation of ADAM22 decreased but did not abolish binding to LGI1 (14). The hydrogen bonds involved in the LGI1_EPTP_–ADAM22_ECD_ interaction might be so weak that the orientation of the hydrogen bonding residues could alter. In the cryo-EM structure, we also found hydrogen bonds between Ser282 of LGI1 and the main-chain carbonyl of Thr397 of ADAM22, between the main-chain carbonyl of Trp376 of LGI1 and Gln334 of ADAM22, and between Lys353 of LGI1 and the main-chain carbonyl of Phe335 of ADAM22 (Fig. S5). These three hydrogen bonds were also formed in the crystal structure.

**Figure 2.**
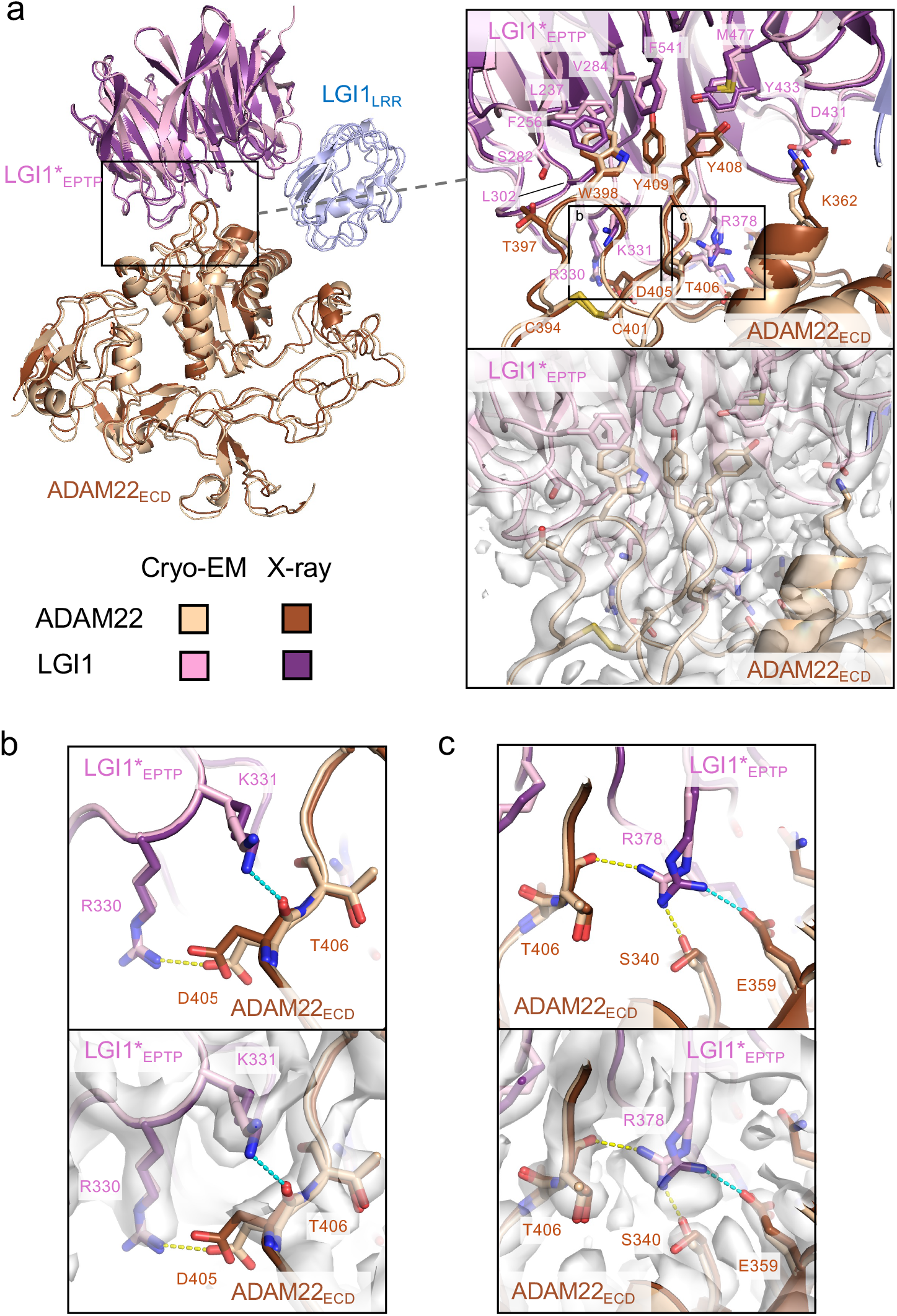
Interactions between LGI1_EPTP_ and ADAM22_ECD_. **a** Overall view of the cryo-EM structure of the LGI1_LRR_–LGI1*_EPTP_–ADAM22_ECD_ complex and a magnified view of the interface between LGI1*_EPTP_ and ADAM22_ECD_. The density map of the interface is shown as white surfaces. The previously reported X-ray structure of the LGI1_EPTP_– ADAM22_ECD_ complex is superposed with distinct colors. Two boxes in the magnified view indicate the locations of the views shown in (**b**; left box) and (**c**; right box). **b** Close-up view of the interaction between Arg330 of LGI1*_EPTP_ and Asp405 of ADAM22_ECD_. The structure (top) and corresponding map (bottom) are shown. Yellow and cyan dashed lines indicate a hydrogen bond observed in the cryo-EM and X-ray structures, respectively. **c** Close-up view of the interaction around Arg378 of LGI1. The structure (top) and corresponding map (bottom) are shown. Yellow and cyan dashed lines indicate hydrogen bonds observed in the cryo-EM and X-ray structures, respectively.

### Intermolecular interactions between LGI1_LRR_ and LGI1*_EPTP_

The present cryo-EM map of the LGI1_LRR_–LGI1*_EPTP_–ADAM22_ECD_ complex provides a more detailed view of the interaction between LGI1_LRR_ and LGI1*_EPTP_ (Fig. 3a) than the previous crystal structure of the 2:2 complex at a moderate resolution of 7.125 Å. Specifically, Glu123 and Arg76 of LGI1_LRR_ form hydrogen bonds with Arg474 and Glu516 of LGI1*_EPTP_, respectively. The hydrogen bond between Glu123 of LGI1_LRR_ and Arg474 of LGI*_EPTP_ is reinforced by stacking with Phe121 of LGI1_LRR_ (Fig. 3a). Additionally, Asn52 and Ser73 of LGI1_LRR_ form hydrogen bonds with the main-chain carbonyls of Tyr496 and Asp495 of LGI1*_EPTP_, respectively. Leu54, Val75, Leu97, and Phe121 of LGI1_LRR_ also form extensive hydrophobic interactions (Fig. 3a). The R474A mutation in LGI1_EPTP_ is a missense ADEAF mutation in LGI1, known to cause epileptic symptoms by inhibiting the assembly of the LGI1–ADAM22 higher-order complex (14, 23, 24). The present higher-resolution structure demonstrates that Arg474 of LGI1*_EPTP_ forms a hydrogen bond with Glu123 of LGI1_LRR_, which was ambiguous in the previous moderate-resolution crystal structure. Regarding interactions for the higher-order assembly, the previous crystal structure suggested a possible interaction between His116 of LGI1_LRR_ and Glu446 of ADAM22 (14). However, in the present cryo-EM structure, the distance between His116 and Glu446 is 6.6 Å (Fig. S6), indicating that they do not interact with each other. No tight interaction was found between LGI1_LRR_ and ADAM22_ECD_ in the cryo-EM structure of the LGI1_LRR_– LGI1*_EPTP_–ADAM22_ECD_ complex.

**Figure 3.**
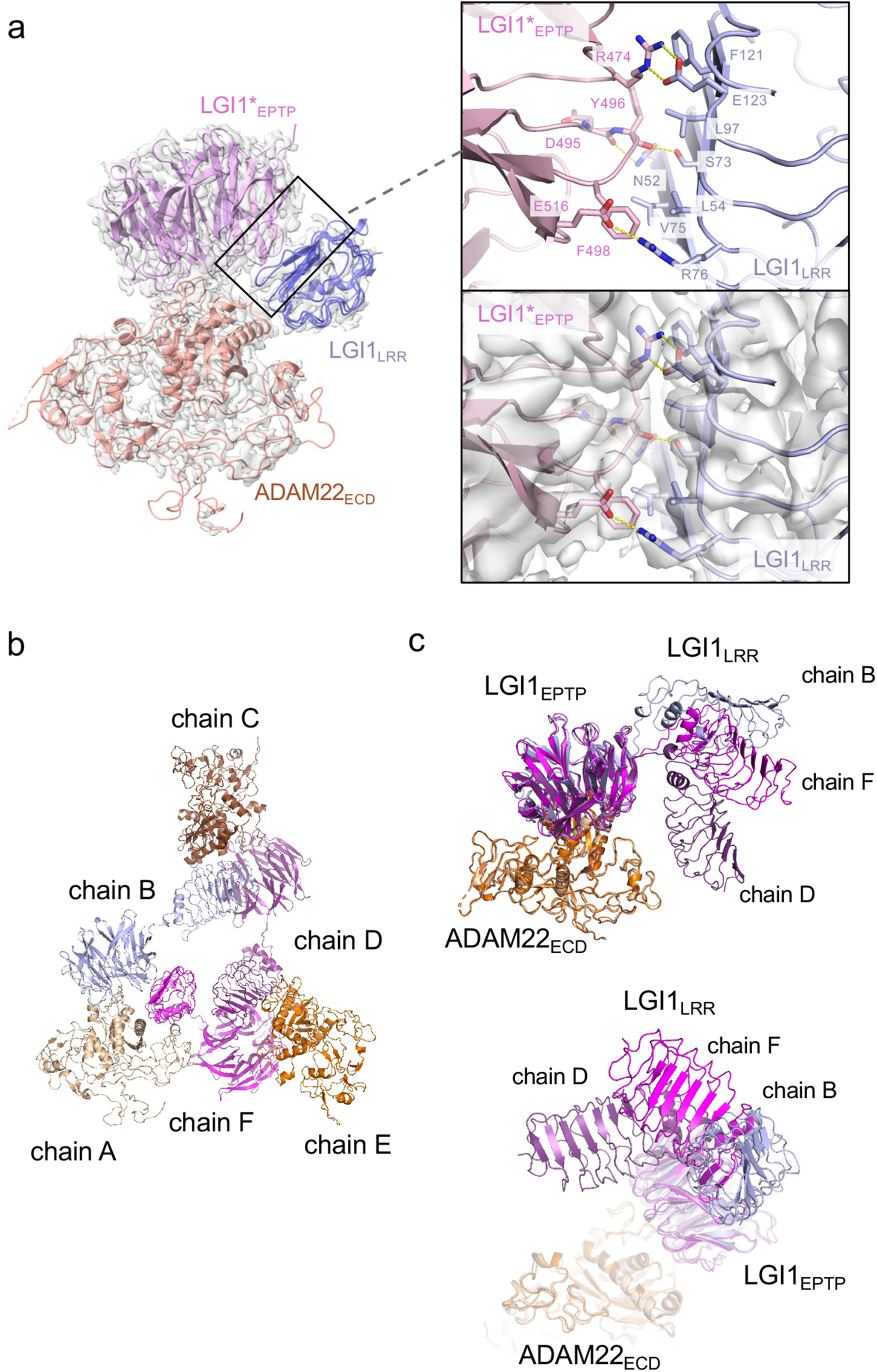
Structure and interaction of the higher-order LGI1–ADAM22_ECD_ complex. **a** Overall view of the cryo-EM structure of the LGI1_LRR_–LGI1*_EPTP_–ADAM22_ECD_ complex and a magnified view of the interface between LGI1_LRR_ and LGI1*_EPTP_. The density map of the interface is shown as white surfaces. **b** Chain IDs of the individual LGI1 or ADAM22_ECD_ molecules in the 3:3 LGI1–ADAM22_ECD_ complex, assigned in this study. **c** Superposition of the three LGI1– ADAM22_ECD_ complexes in the cryo-EM structure of the 3:3 LGI1-ADAM22_ECD_ complex, using LGI1*_EPTP_–ADAM22_ECD_ as the reference.

### Cryo-EM structure of the 3:3 LGI1–ADAM22_ECD_ complex

The cryo-EM structure of the LGI1_LRR_–LGI1*_EPTP_–ADAM22_ECD_ complex could fit a density map of the 3:3 LGI1–ADAM22_ECD_ complex well. Three LGI1_LRR_–LGI1*_EPTP_–ADAM22_ECD_ complex structures were put into the 3.79-Å-resolution map of the 3:3 complex. The density of LGI1 and the metalloprotease-like domain of ADAM22_ECD_ appears relatively strong. The density of one interface between LGI1*_EPTP_ and ADAM22_ECD_ is especially resolved well (Fig. S7; see the interface between chains A and B). Although the density of amino-acid side chains in LGI1_LRR_ is not well separated, the main-chain structure of LGI1_LRR_ could be fitted to the density map (Fig. S7; see the density of LGI1_LRR_ in chain F). On the other hand, the density of the disintegrin, cysteine-rich, and EGF-like domains after Pro445 of ADAM22 is relatively weak in all three molecules. Correspondingly, in the local resolution map of the 3:3 LGI1–ADAM22_ECD_ complex, the resolution of these ADAM22 domains after Pro445 is low (Fig. S8). Like the crystal structure of the 2:2 LGI1–ADAM22_ECD_ complex, the LRR and EPTP domains of LGI1 are linked by a two-residue linker (Ile222-Ile223) in an extended conformation. LGI1_EPTP_ interacts with the metalloprotease-like domain of ADAM22 to form the LGI1_EPTP_–ADAM22_ECD_ complex, while the LRR domain of one LGI1 molecule interacts with the EPTP domain of the neighboring LGI1, thereby bridging three distant ADAM22 molecules in the 3:3 complex (Fig. 1d,e). The C-termini of the three ADAM22 molecules point in opposite directions. When the three assembled LGI1– ADAM22_ECD_ complexes were superposed with the LGI1_EPTP_–ADAM22_ECD_ structure as the reference, all three LGI1 LRR domains were oriented differently (Fig. 3b,c). Although the triangular shape observed in the 2D class averaged image suggested (pseudo-)*C*3 symmetry of the 3:3 complex, the determined structure of the 3:3 complex was not symmetric. Actually, the *C*3 symmetry-restrained 3:3 model we previously calculated based on the SEC-SAXS analysis using the program SASREF (14, 25) could not be fitted with the present cryo-EM structure (Fig. S9a,b). This discrepancy arises from the difference in the orientation of LGI1_LRR_ relative to the LGI1*_EPTP_–ADAM22 complex between the previous model and the present cryo-EM structure (indicated by the arrowhead in Fig. S9b). The structure of the 3:3 LGI1–ADAM22 complex was also predicted by AlphaFold3 (26), which suggested a *C*3 symmetric assembly (Fig. S9a). Intriguingly, the orientation of LGI1_LRR_ relative to LGI1*_EPTP_–ADAM22 was similar to that observed in the present cryo-EM structure (indicated by the arrowhead in Fig. S9c), despite relatively low prediction accuracy of the assembly (ipTM = 0.47, pTM = 0.52; pLDDT color outputs and PAE plot are shown in Fig. S9d,e). On the other hand, the overall trimeric configuration is different between the *C*3 symmetric AlphaFold3 model and the non-symmetric cryo-EM structure (Fig. S9c).

We then analyzed the interdomain motion of the three LGI1 domains (assigned to chains B, D, and F in the deposited PDB file; Fig. 3b) in the 3:3 LGI1–ADAM22_ECD_ complex by the DynDom server (27) (Fig. S10 and Table S2). Analysis of the motion between the three pairs of chains suggested that the static domain corresponds to the EPTP domain, while the mobile domain corresponds to the LRR domain, predicting a hinge region for interdomain bending. The LRR domain of chain D was rotated 69.8° around the hinge axis compared to chain B, and the LRR domain of chain F was rotated 69.0° around the hinge axis compared to chain B. The LRR domains of chain D and chain F were also rotated 70.0° around the hinge axis relative to each other, suggesting that each LRR domain rotates approximately 70° about the hinge axis in the 3:3 LGI1–ADAM22_ECD_ complex (Table S2). Furthermore, the motion of each LRR domain varied widely: The movement of the LRR domain of chain D with respect to chain B follows a standard closure motion of 14 %, whereas the movement of the LRR domain of chain F with respect to

chain D follows a standard closure motion of 99 % (Table S2). This closure motion appears to locate the LRR domain of chain F in close proximity to that of chain D to make the triangular assembly slightly more compact, which might stabilize the non-symmetric trimeric configuration observed in the cryo-EM structure.

### Dynamics of the LGI1−ADAM22 higher-order complex observed by HS-AFM

To directly visualize the molecular dynamics of the LGI1–ADAM22_ECD_ complex and characterize its structural properties in solution, we performed HS-AFM (21, 22). HS-AFM images of gel-filtration chromatography fractions containing the 3:3 LGI1–ADAM22_ECD_ complex (not chemically crosslinked with glutaraldehyde) predominantly revealed triangular-shaped molecules, which appeared to exist stably with no drastic structural changes (Fig. 4a-c and Movie S1). A comparison with the simulated AFM image suggests that the protrusion on the exterior of the triangle is likely ADAM22 (Fig. 4b). This site frequently appeared to dissociate during HS-AFM scanning (Fig. 4c and Movie S1), indicating that the interaction between LGI1 and ADAM22 is weaker than the interactions among LGI1 molecules within the 3:3 LGI1–ADAM22_ECD_ assembly. Together with the cryo-EM structure, this also indicates that the trimerization can be entirely organized by LGI1, suggesting the possibility that LGI1 could trimerize on its own, although this possibility could not be tested due to the difficulty in the expression of the full-length LGI1 alone for biophysical analysis in our hands. On the other hand, considering the dynamic property of the 3:3 complex and spatial alignment of LGI1_LRR_ and ADAM22_ECD_, we cannot exclude the possibility that ADAM22 could act as a platform to facilitate the intermolecular interaction between LGI1_LRR_ and LGI1*_EPTP_ for the trimerization of LGI1.

**Figure 4.**
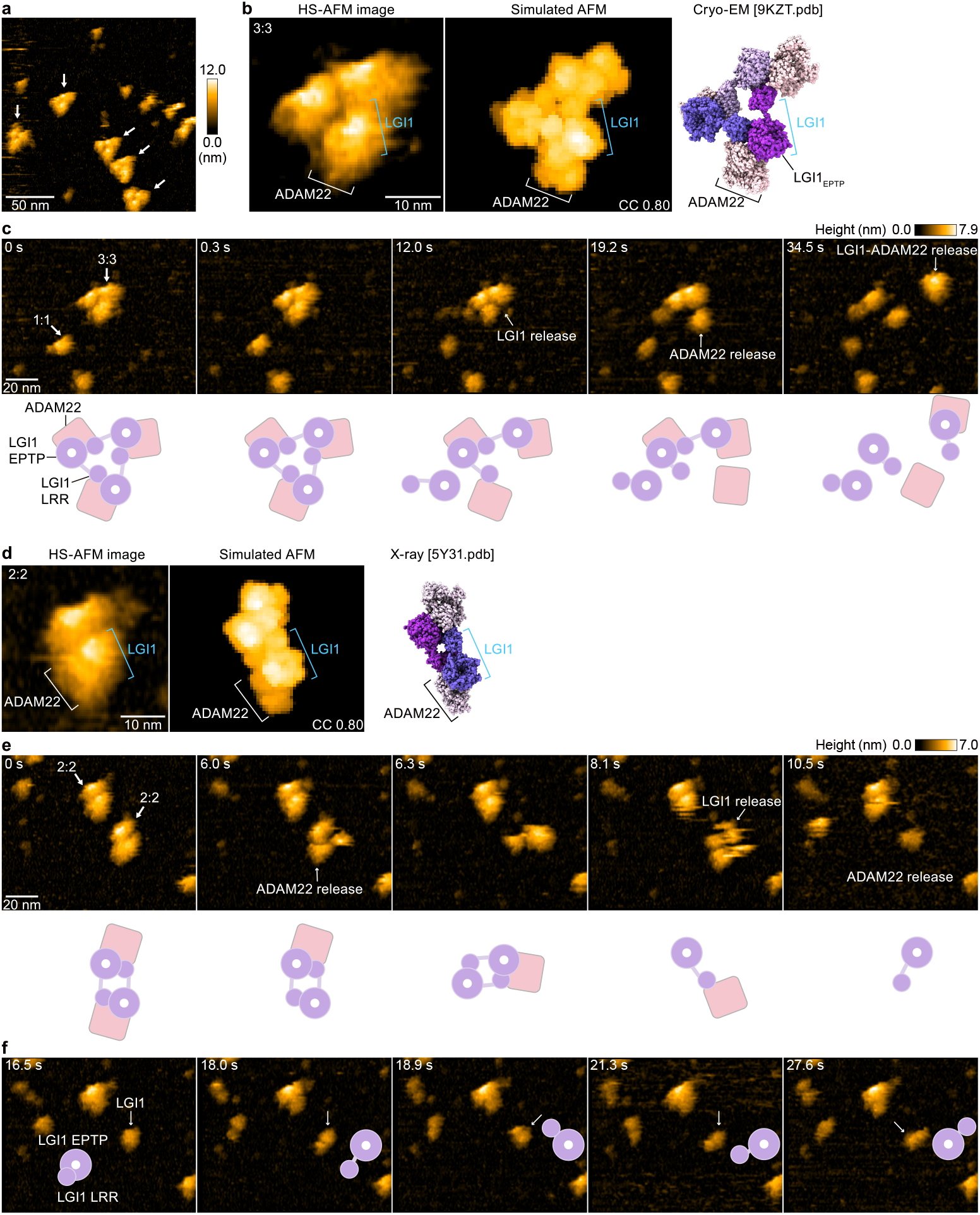
HS-AFM observations of LGI1–ADAM22_ECD_ complexes. **a** A representative HS-AFM image of the 3:3 LGI1–ADAM22_ECD_ complex. The color bars on the right indicate height in nanometers. White arrows indicate the 3:3 complex. The frame rate was 1.0 frames/s. **b, d** Magnified HS-AFM images of the 3:3 (left in **b**) and 2:2 (left in **d**) LGI1–ADAM22_ECD_ complexes. The simulated AFM images (middle) were derived from fitting to the experimental HS-AFM image (left). The well-fitting simulated AFM images and the coefficient of correlation (CC) are indicated. The cryo-EM structure of the 3:3 complex (right in **b**) and the X-ray structure of the 2:2 complex (right in **d**) are shown in the same orientation as the simulated AFM images. **c, e, f** Sequential HS-AFM images of the 3:3 (c; see also Movie S1) and 2:2 (**e, f**; see also Movie S2) LGI1–ADAM22_ECD_ complexes. A schematic illustration of the interpretation of HS-AFM images is shown at the bottom. Imaging parameters: scanning area = 120 × 96 nm^2^ (240 × 192 pixels); frame rate = 3.3 frames/s. HS-AFM experiments were repeated independently at least 3 times with consistent results.

In addition to the 3:3 complex, HS-AFM also captured the dynamics of the 2:2 LGI1-ADAM22_ECD_ complex (Fig. 4d,e and Movie S2). A comparison of the experimental HS-AFM image with the simulated AFM image based on the crystal structure of the 2:2 complex indicates that the outer site corresponds to ADAM22, suggesting that LGI1 and ADAM22 are facing each other. The 2:2 complex also exhibited fragility during HS-AFM imaging, similar to the 3:3 complex (Fig. 4e and Movie S2). In individual LGI1 molecules dissociated from the complex, the LRR domain moved freely relative to the EPTP domain (Fig. 4f and after 10.2 s in Movie S2), likely reflecting the conformational difference in the relative orientation between LGI1_LRR_ and LGI1_EPTP_ observed within the cryo-EM structure of the 3:3 complex. Thus, single-molecule observations using HS-AFM demonstrated that the 3:3 LGI1–ADAM22_ECD_ complex is present in solution, that the binding between ADAM22 and LGI1 is relatively weak within the 3:3 LGI1– ADAM22_ECD_ assembly, and that the LRR domain of LGI1 exhibits flexibility.

## Discussion

In this study, the ligand–receptor complex between LGI1, a secreted protein of neurons, and its receptor protein, ADAM22, was investigated by cryo-EM and HS-AFM. By chemically crosslinking with glutaraldehyde, we successfully captured much larger numbers of particle images of the 3:3 LGI1–ADAM22_ECD_ complex than in our initial, preliminary cryo-EM study (14). We could determine not only the overall 3D structure of the 3:3 LGI1–ADAM22_ECD_ complex but also the 3D structure of the LGI1_LRR_–LGI1*_EPTP_–ADAM22_ECD_ complex at a nominal resolution of 2.78 Å, revealing the interaction mode of the LGI1–ADAM22 higher-order complex at a higher resolution than before. HS-AFM successfully imaged the LGI1–ADAM22_ECD_ higher-order complex and confirmed that LGI1–ADAM22_ECD_ forms a 3:3 complex in solution.

The present 3.79-Å-resolution cryo-EM map of the 3:3 LGI1–ADAM22_ECD_ complex was calculated from 120,708 particle images selected after two rounds of heterogeneous refinement. On the other hand, a few classes of other particle images display triangular shapes with missing parts, suggesting domain motions or conformational heterogeneity in the 3:3 complex (Fig. S4). This raised a possibility that motion-based refinement might improve resolution in flexible regions. Therefore, we first performed 3D flexible refinement (3D Flex) using CryoSPARC (Movie S3) (28). However, even with the 3D Flex refinement, the density of the disintegrin, cysteine-rich, and EGF-like domains in the ADAM22 molecule remained poorly resolved. Then, we next hypothesized that the 3:3 LGI1–ADAM22_ECD_ complex undergoes discrete, non-rigid motions and performed 3D Variability Analysis (3D VA) in CryoSPARC (29). Using three orthogonal principal modes, the 3D VA analysis indicated two twisting motions and one stretching motion of the triangular-shaped 3:3 LGI1–ADAM22_ECD_ complex (Movie S4). This analysis also visualized relatively large motions of the disintegrin, cysteine-rich, and EGF-like domains of ADAM22 (Movie S4), suggestive of intrinsic conformational flexibilities of these domains, likely resulting in lower local resolution in the periphery of the complex (Fig. S8).

Previous studies have shown that LGI1 is enriched not only at the synapse but also at the axon initial segment and colocalized with ADAM22/23 and the voltage-gated potassium (Kv1) channels (18, 20). The Kv1-associated cell-adhesion molecules, TAG-1 and Caspr2, likely mediate this colocalization by binding to ADAM22/23. LGI1 knock-out mice show a reduced density of Kv1 channels, which is associated with increased neuronal excitability of hippocampal CA3 neurons, indicating that LGI1 regulates Kv1 channel function (18). In addition, the LGI1 R474Q mutation has been found to interfere with the colocalization of ADAM22/23 and Kv1 channels (20). Given that the LGI1 R474Q mutation inhibits the higher-order assembly of LGI1– ADAM22 with little impact on LGI1 secretion and binding to ADAM22, the higher-order complex of LGI1–ADAM22 likely regulates the axonal Kv1 channel function. The 3:3 LGI1– ADAM22 complex observed *in vitro* might facilitate efficient clustering of the axonal Kv1 channels to control their density and inhibit epilepsy. A recent study has shown that LGI3, a member of the LGI family, selectively co-assembles with Kv1 channels by using ADAM23 as the receptor in axons (30). LGI3 is secreted by oligodendrocytes in the brain and enriched at juxtaparanodes of myelinated axons to form subclusters. LGI1 and LGI3, along with ADAM22 and ADAM23, belong to the same subfamily and share a similar domain organization. This suggests that the LGI3–ADAM23 complex, like the LGI1–ADAM22 complex, may form a higher-order assembly, which could be related to the clustering mechanism of Kv1 channels in axons. In this context, as discussed in (30), either or both of the 2:2 and 3:3 complexes might be formed in a *trans* fashion at the juxtaparanode of myelinated axons and bridge the axon and the innermost myelin membrane. Alternatively, the 3:3 complex formed in a *cis* fashion might positively regulate the clustering of the axonal Kv channels at the juxtaparanode, possibly in a similar manner at the axon initial segment. Finally, this study proposes that the LGI1–ADAM22 complex is an interesting therapeutic target for epilepsy and other neurological disorders. The 3:3 LGI1–ADAM22 complex structure revealed in this study could serve as a useful platform for structure-based drug design and facilitate the development of antiepileptic drugs.

## Materials and methods

### Protein preparation

For preparation of the LGI1–ADAM22_ECD_ complex, the C-terminally His_6_-tagged LGI1 (R470A) was co-expressed with the non-tagged ADAM22_ECD_ in Expi293F cells (Thermo Fisher Scientific). As reported previously, the R470A mutation of LGI1 increases the yield of the LGI1– ADAM22_ECD_ complex without affecting the higher-order assembly of LGI1–ADAM22_ECD_ (14). The vectors for the co-expression were the pEBMulti-Neo vector (Wako chemicals) harboring the gene encoding human LGI1 (full length, residues 37–557; R470A) with the N-terminal Igκ signal sequence and that harboring the gene encoding human ADAM22_ECD_ including the N-terminal prosequence (residues 1–729), both of which were reported previously (14). The culture media were loaded onto a Ni-NTA (Qiagen) column pre-equilibrated with 20 mM Tris-HCl (pH 8.0) containing 300 mM NaCl. After the column was washed with 20 mM Tris-HCl (pH 8.0) containing 300 mM NaCl and 25 mM imidazole, the proteins were eluted with 20 mM Tris-HCl (pH 8.0) containing 300 mM NaCl and 250 mM imidazole. The eluted proteins were further purified by size-exclusion chromatography using Superdex200 (GE healthcare) with 20 mM Tris-HCl (pH 7.5) buffer containing 50 mM NaCl. The purified proteins were concentrated to 0.25 g/L in Amicon Ultra-15 50,000 MWCO filter (Millipore), and glutaraldehyde was added at a final concentration of 0.1% (v/v). The sample was concentrated to 500 µL in Amicon Ultra-4 50,000 MWCO filter (Millipore) and purified by gel filtration chromatography using a Superose6 10/300 GL (GE Healthcare) column with 20 mM Tris-HCl buffer (pH 7.5) containing 50 mM NaCl. One of the peak fraction was concentrated to 0.5 g/L by Amicon Ultra-0.5 50,000 MWCO filter (Millipore).

### Cryo-EM single particle analysis

Cu grids (R1.2/1.3, 300 mesh, Quantifoil) with holey carbon films were hydrophilized using a JEC-3000FC Auto Fine Coater (JEOL) at 7 Pa, 10 mA, and 10 sec. A 3 μL aliquot of 0.5 g/L protein solution was added to the grids using a Vitrobot Mark IV (Thermo Fisher Scientific) at a temperature of 8°C and 100% humidity. The grids were then immersed in liquid ethane and rapidly frozen under the following conditions: waiting time of 0 sec, blotting time of 3 sec, and blotting force of 10. Data collection were carried out on a CRYO ARM 300 transmission electron microscope (JEOL Ltd., Japan) operating at 300 kV, equipped with an Omega-type in-column energy filter (slit width 20 eV) and a Gatan K3 electron detector (operated in correlated doubling sample mode) at SPring-8 (Hyogo, Japan). A total of 7,625 movies were automatically collected using SerialEM (31). Hole centering was performed using yoneoLocr (32) integrated as a SerialEM macro. Movies were collected using the beam-image shift method (5×5×1 matrices), at a target defocus range of −1.4 to −1.7 μm and a nominal magnification of ×60,000, corresponding to a calibrated pixel size of 0.752 Å/pixel. Each movie was recorded with an exposure time of 2.79627 s, subdivided into 60 frames with a total electron dose of 60.8046 e^−1^ Å^−2^.

All processing was performed in CryoSPARC v.4.1.2 or higher (33). The collected micrographs were processed using patch motion correction and patch CTF correction. Particles were picked from 7,635 particle images with using a template, yielding 3,336,559 particles. After extracting the particles from the micrographs and classifying them into 2D classes, about 60% of the particles based on the average 2D image of the protein were selected. An initial 3D reconstruction and heterogeneous refinement were performed on six classes, and one of these classes was further refined by non-uniform refinement to obtain a 2.78 Å resolution 3D map of the LGI1_LRR_–LGI1*_EPTP_–ADAM22_ECD_ complex. Additionally, by performing *ab initio* reconstruction and heterogeneous refinement of one of the six classes into three subclasses and refining one of these subclasses further with non-uniform refinement (34), a 3D map of the 3:3 LGI1–ADAM22_ECD_ complex with a resolution of 3.79 Å was obtained. Overall resolution estimates correspond to an Fourier shell correlation of 0.143 using an optimized mask that is automatically determined after refinement. Local resolution maps were obtained using local resolution estimation. Movies were created using 3D VA (29) and 3D Flex (28).

### Model building

Model building was performed using the programs Coot (35) and UCSF ChimeraX (36). The initial model of the LGI1_LRR_–LGI1*_EPTP_–ADAM22_ECD_ complex was built using a part of the crystal structure of the 2:2 LGI1–ADAM22_ECD_ complex (PDB 5Y31) (14) and fitted into the cryo-EM map. The initial model of the 3:3 LGI1–ADAM22_ECD_ complex was built using the cryo-EM structure of the LGI1_LRR_–LGI1*_EPTP_–ADAM22_ECD_ complex. Three LGI1_LRR_–LGI1*_EPTP_– ADAM22_ECD_ complexes were fitted into the cryo-EM map and LGI1_LRR_ and LGI1_EPTP_ were connected by modeling the linker region. The structure refinement was performed using the Phenix software package (37). All figures and movies were created using the programs PyMOL (38) and UCSF ChimeraX (36).

### HS-AFM observations

HS-AFM experiments were conducted using a custom-built high-speed atomic force microscope operating in tapping mode (22). Briefly, an optical beam deflection detector monitored the cantilever’s (BL-AC10DS-A2, Olympus, Japan) deflection using a 780 nm, 0.8 mW infrared (IR) laser. The cantilever exhibited a spring constant of approximately 100 pN/nm, a resonant frequency near 400 kHz, and a quality factor around 2 in a liquid environment. The IR beam was directed onto the cantilever’s back surface through a 60× objective lens (CFI S Plan Fluor ELWD 60X, Nikon, Japan), and its reflection was detected by a two-segmented PIN photodiode (MPR-1, Graviton, Japan). The initial AFM tip had a triangular section resembling a bird’s beak. To enhance spatial resolution, an amorphous carbon tip was grown on the bird beak tip via electron beam deposition using a scanning electron microscope (FE-SEM; Verious 5UC, Thermo Fisher Scientific, USA). The additional AFM tip was roughly 500 nm long, with an apex radius of approximately 1 nm after plasma etching with a plasma cleaner (Tergeo, PIC Scientific, USA). All HS-AFM images were obtained using the cantilever with the additional AFM tip. The cantilever’s free oscillation amplitude was less than 1 nm, and during HS-AFM scanning, the set-point amplitude was adjusted to roughly 90% of the free amplitude. To minimize interaction forces between the sample and the AFM tip, “only trace imaging” (OTI) mode (39) was employed for HS-AFM. Data acquisition and operation were managed using custom software based on Visual Basic. NET (Microsoft).

HS-AFM observations of the LGI1–ADAM22_ECD_ complex (not chemically crosslinked with glutaraldehyde) were conducted on AP-mica, prepared by treating the mica surface with 0.00005% (3-aminopropyl)triethoxysilane (APTES) (Shin-Etsu Chemical, Japan) in MilliQ water for 3 min. Samples at a concentration of 10 nM were added to the AP-mica surface after 3 min incubation of 3 µL. HS-AFM observations of the LGI1–ADAM22_ECD_ complex were performed in 20 mM Tris-HCl (pH 7.4) buffer containing 150 mM NaCl. All HS-AFM experiments were performed at room temperature (24–26°C) and was independently repeated at least three times, consistently yielding similar results. For image processing, the HS-AFM images were processed using Fiji (ImageJ) software (NIH, USA) (40). A mean filter with a radius of 0.5 pixels was utilized to lower noise levels in each image. The Template Matching and Slice Alignment plugin for ImageJ was used to correct for drift between sequential images.

### Simulation of AFM images

The BioAFMviewer software (41) was utilized to validate the captured topographies of the LGI1– ADAM22_ECD_ complex. The simulated scanning was based on the nonelastic collisions between a rigid cone-shaped tip model and the rigid van der Waals atomic model of the protein structure . Automatized fitting (42) was employed to generate a simulated image that closely matched the HS-AFM target image (image correlation coefficients reported in each figure). In the simulation presented in Fig. 4, the tip shape parameters were set to *R* = 0.4 nm for the tip probe sphere radius and *α* = 5.0° for the cone half angle.

## Data, materials, and software availability

The coordinates and maps of the LGI1_LRR_–LGI1*_EPTP_–ADAM22_ECD_ complex and the 3:3 LGI1– ADAM22_ECD_ complex have been deposited in the Protein Data Bank/Electron Microscopy Data Bank under the accession codes of 9KZC/EMD-62659 and 9KZT/EMD-62668, respectively. Other data are available from the corresponding authors upon reasonable request.

## Acknowledgements

This research was partially supported by Research Support Project for Life Science and Drug Discovery (Basis for Supporting Innovative Drug Discovery and Life Science Research (BINDS)) from AMED under Grant Number JP24ama121001. Preliminary cryo-EM experiments were performed at EM01CT/EM02CT with the approval of the Japan Synchrotron Radiation Research Institute (JASRI) (Proposal No. 2021B2542, 2022A2542, 2022B2543, and 2023A2543). This work was supported by JSPS/MEXT KAKENHI (19H03162 to A.Y., 23H00374 to M.F., 18H03983 and 23H04070 to S.F.), Japan Agency for Medical Research and Development (JP24wm0625319 to Y.F. and JP24ek0109649 to M.F.), Takeda Science Foundation (to S.F.), the World Premier International Research Center Initiative (WPI), MEXT, Japan (to M.S.).

## Author contributions

T.Y. prepared the sample for cryo-EM. T.Y., M.K., and A.M. performed cryo-EM analysis. K.O., A.Y., and S.F. assisted with cryo-EM analysis. A.M. prepared the sample for HS-AFM. M.S. performed HS-AFM experiments and analysis. Y.F., M.F., M.S., and S.F. designed the experiments. A.M., M.S., and S.F. wrote the paper. All authors discussed the results and commented on the manuscript.

## Competing interests

The authors declare no competing interest.

## Notes

### Competing Interest Statement

The authors have declared no competing interest.

### Summary of Updates

An error in the Materials and methods was corrected.

