## Supporting Information for "Structural insights into heterohexameric assembly of epilepsy-related ligand–receptor complex LGI1–ADAM22"

#### Figure legends of Figures S1-S10 and Movies S1-S4

**Figure S1. Purification of the 3:3 LGI1–ADAM22$_{ECD}$ complex.**

**a, b** Gel filtration chromatography (**a**) and SDS-PAGE (**b**) of the 3:3 LGI1–ADAM22$_{ECD}$ complex before crosslinking. The red arrow indicates the fractions subjected to SDS-PAGE. The orange boxes represent the fractions used for crosslinking. **c, d** Gel filtration chromatography (**c**) and SDS-PAGE (**d**) of the 3:3 LGI1–ADAM22$_{ECD}$ complex after crosslinking. The red arrow indicates the fractions subjected to SDS-PAGE. The orange boxes represent the fractions used for cryo-EM analysis.

**Figure S2. 2D class averages of the particles from blob picker in cryoSPARC.**

1,403,037 particles were picked without templates and classified into 100 classes. The particle images surrounded by magenta boxes were selected as templates for template picking.

**Figure S3. 2D class averages of the particles from template picker in cryoSPARC.**

2,530,790 particles were picked with templates and classified into 100 classes. The particle images surrounded by magenta boxes were subjected to *ab initio* reconstruction, whereas those surrounded by cyan boxes were excluded.

**Figure S4. Cryo-EM data and processing for the LGI1$_{LRR}$–LGI1*$_{EPTP}$–ADAM22$_{ECD}$ and 3:3 LGI1–ADAM22$_{ECD}$ complexes.**

Representative 2D class averages are shown in the "Select 2D classes". The obtained final maps are shown as white surfaces with protein structures. The red rounded boxes indicate the density maps that were supposed to correspond to the 3:3 LGI1–ADAM22$_{ECD}$ complex but were not used to calculate the final map.

**Figure S5. Comparison of the LGI1$_{EPTP}$–ADAM22$_{ECD}$ interaction between the cryo-EM and crystal structures.**

The present cryo-EM structure of the LGI1$_{LRR}$–LGI1*$_{EPTP}$–ADAM22$_{ECD}$ complex and the previous X-ray crystal structure of the LGI1$_{EPTP}$–ADAM22$_{ECD}$ complex are superposed. Close-up views (bottom panels) highlight amino acid residues involved in hydrogen bonds between LGI1$_{EPTP}$ and ADAM22$_{ECD}$ that were observed in the cryo-EM structure but not in the previous X-ray structure. These hydrogen bonds are indicated by dashed yellow lines. Density maps are shown as white surfaces.

**Figure S6. LGI1$_{LRR}$ and ADAM22$_{ECD}$ in the LGI1–ADAM22$_{ECD}$ assembly.**

LGI1$_{LRR}$ is located in close proximity to ADAM22$_{ECD}$. However, no tight interaction was observed between LGI1$_{LRR}$ and ADAM22$_{ECD}$ in the cryo-EM structure.

**Figure S7. Cryo-EM density map of the 3:3 LGI1-ADAM22$_{ECD}$ complex.**

Density maps of $LGI1_{LRR}$ in chain F, $LGI1_{EPTP}$ in chain B, and the interface between $LGI1_{EPTP}$ in chain B and $ADAM22_{ECD}$ in chain A are shown, along with the overall view of the map of the 3:3 LGI1–$ADAM22_{ECD}$ complex. The labeled residues are important for the $LGI1_{EPTP}$–$ADAM22_{ECD}$ interaction.

**Figure S8. Local resolution of the cryo-EM structure of the 3:3 LGI1–$ADAM22_{ECD}$ complex.**
Distribution of the resolution was plotted onto the density at high (0.06) and low (0.03) levels.

**Figure S9. Comparison of the 3:3 LGI1–$ADAM22_{ECD}$ assembly determined by cryo-EM with that calculated based on SEC-SAXS analysis and that predicted by AlphaFold3.**
**a** SEC-SAXS-based model (left), cryo-EM structure (middle), and AlphaFold3-predicted model (right). **b** Superposition of the SEC-SAXS-based model with *C*3 symmetry restraints and the cryo-EM structure. The structures were aligned based on $LGI1_{EPTP}$ in chain A from the cryo-EM structure. The orientation of $LGI1_{LRR}$ (indicated by the arrowhead) differs between these two structures. **c** Superposition of the *C*3 symmetric model predicted by AlphaFold3 and the cryo-EM structure. The structures were aligned based on $LGI1_{EPTP}$ in chain A from the cryo-EM structure. The orientation of $LGI1_{LRR}$ (indicated by the arrowhead) is similar between these two structures although the overall trimeric configuration is different. **d** AlphaFold3-predicted model colored according to pLDDT confidence scores (orange: 0–50; yellow: 50–70; cyan: 70–90; blue: 90–100) **e** Predicted Aligned Error (PAE) matrix of the AlphaFold3 prediction.

**Figure S10. Conformational difference of the three LGI1 molecules in the 3:3 LGI1–$ADAM22_{ECD}$ assembly.**
The conformational difference was analyzed by the DynDom server. The superpositions of chains B and D (**a**), chains B and F (**b**), and chains D and F (**c**), along with their rotations about the hinge axis, are shown individually. Black arrows indicate the rotations of $LGI1_{LRR}$ relative to $LGI1_{EPTP}$.

**Movie S1. HS-AFM videos of three representative 3:3 LGI1–$ADAM22_{ECD}$ complexes on the AP-mica, related to Figure 4c.**
White, blue, and magenta arrows indicate LGI1 release, ADAM22 release and LGI1–ADAM22 release, respectively. Image size, 240 × 192 $pixels^2$; scan area, 120 × 96 $nm^2$; frame rate, 3.3 fps.

**Movie S2. HS-AFM videos of a representative 2:2 LGI1–$ADAM22_{ECD}$ complexes on the AP-mica, related to Figure 4e.**
White and blue arrows indicate LGI1 release and ADAM22 release, respectively. Image size, 240 × 192 $pixels^2$; scan area, 120 × 96 $nm^2$; frame rate, 3.3 fps.

**Movie S3. 3D Flex analysis of the 3:3 LGI1–ADAM22$_{ECD}$ complex.**

Two latent coordinates were used in the 3D Flex analysis. Two 3D Flex movies (**a**, **b**) are displayed at the same threshold.

**Movie S4. 3D VA analysis of the 3:3 LGI1–ADAM22$_{ECD}$ complex.**

Three variability components are represented in the 3D VA analysis. Three 3D VA movies (**a**-**c**) are displayed at the same threshold. Two movies (**a**, **b**) show twisting motion, whereas one movie (**c**) shows stretching motion.

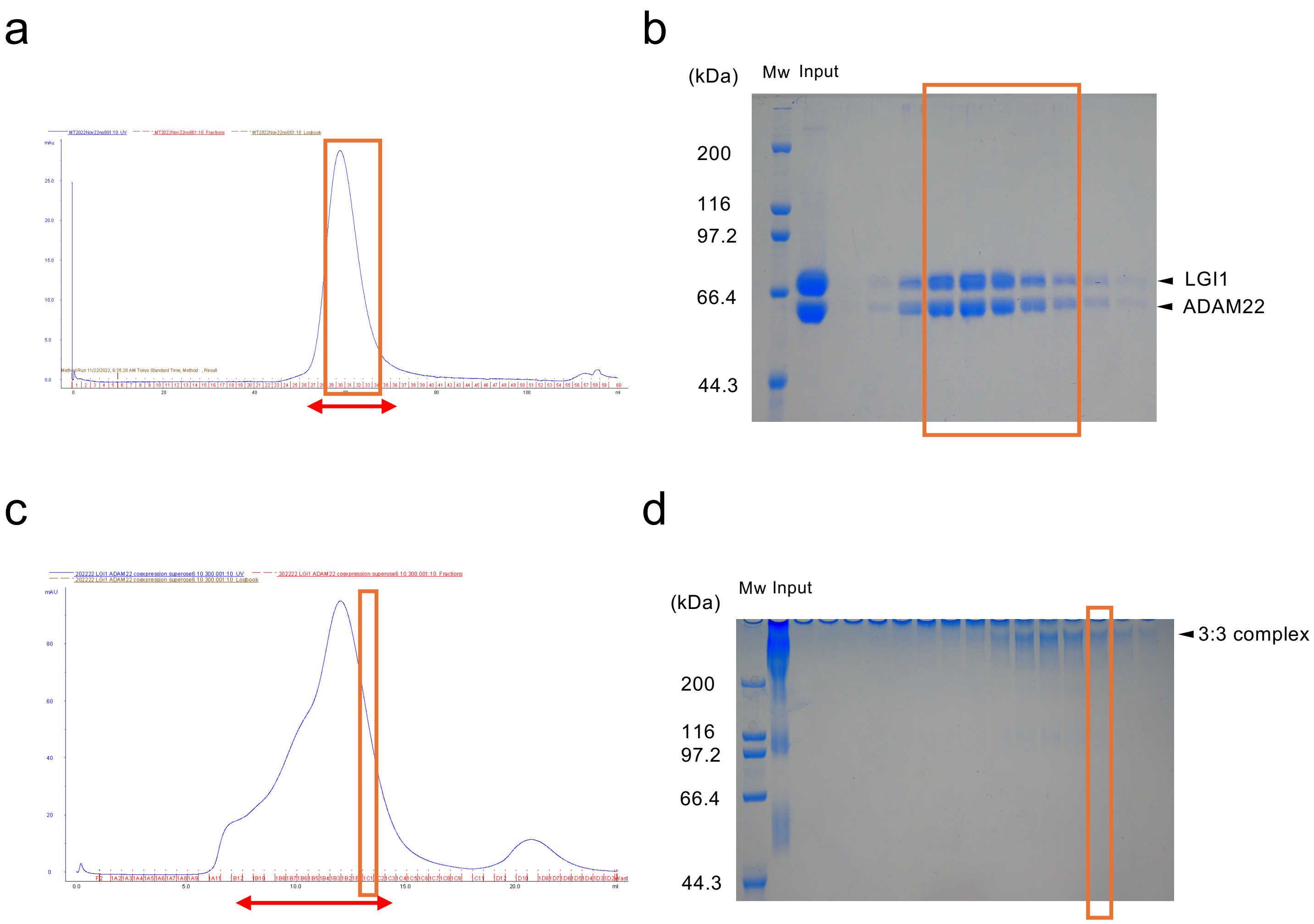
a
b
(kDa)
Mw
Input
200
116
97.2
66.4
44.3
LGI1
ADAM22
c
d
(kDa)
Mw
Input
3:3 complex
200
116
97.2
66.4
44.3

Fig. S1

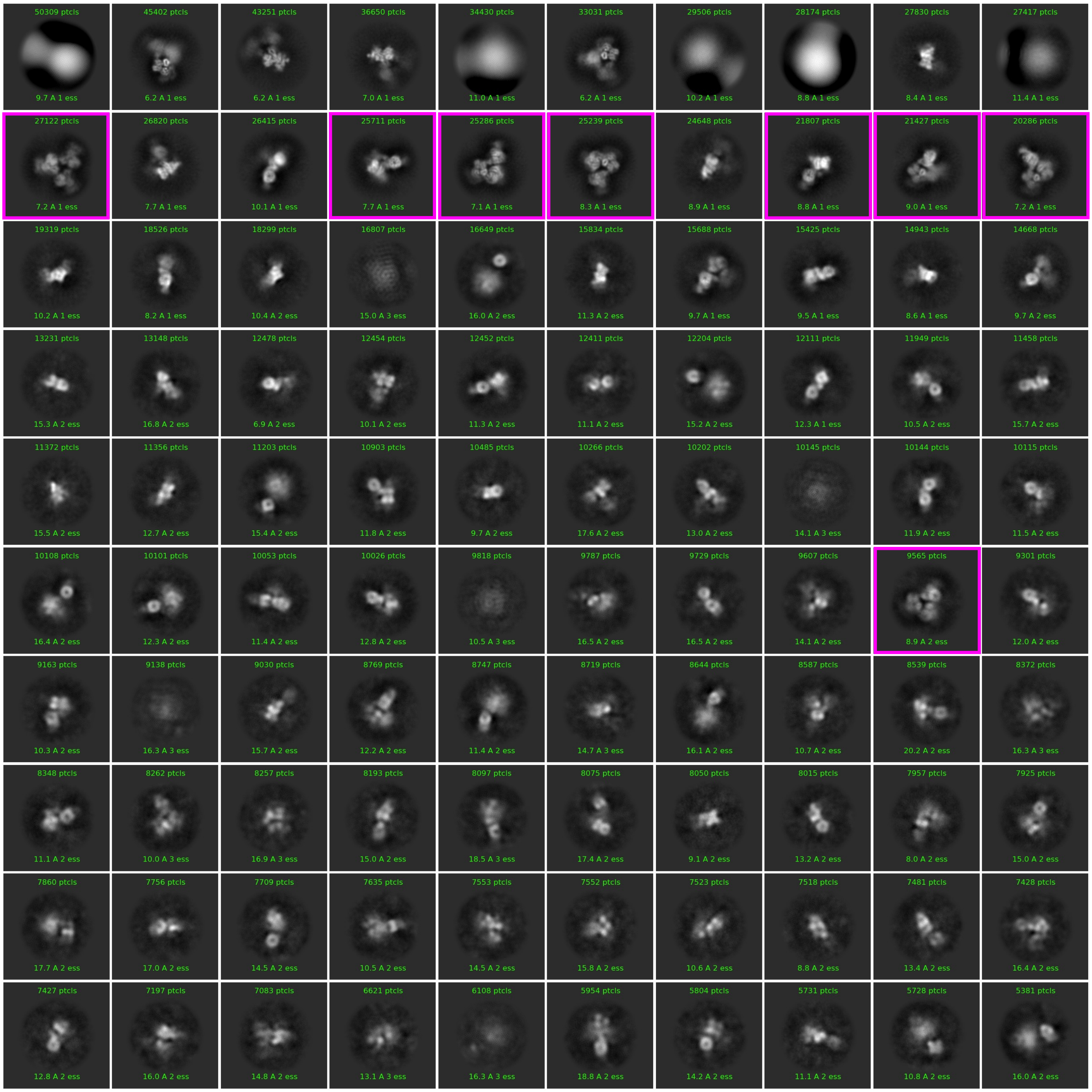
50309 ptcls
9.7 A 1 ess
45402 ptcls
6.2 A 1 ess
43251 ptcls
6.2 A 1 ess
36650 ptcls
7.0 A 1 ess
34430 ptcls
11.0 A 1 ess
33031 ptcls
6.2 A 1 ess
29506 ptcls
10.2 A 1 ess
28174 ptcls
8.8 A 1 ess
27830 ptcls
8.4 A 1 ess
27417 ptcls
11.4 A 1 ess
27122 ptcls
7.2 A 1 ess
26820 ptcls
7.7 A 1 ess
26415 ptcls
10.1 A 1 ess
25711 ptcls
7.7 A 1 ess
25286 ptcls
7.1 A 1 ess
25239 ptcls
8.3 A 1 ess
24648 ptcls
8.9 A 1 ess
21807 ptcls
8.8 A 1 ess
21427 ptcls
9.0 A 1 ess
20286 ptcls
7.2 A 1 ess
19319 ptcls
10.2 A 1 ess
18526 ptcls
8.2 A 1 ess
18299 ptcls
10.4 A 2 ess
16807 ptcls
15.0 A 3 ess
16649 ptcls
16.0 A 2 ess
15834 ptcls
11.3 A 2 ess
15688 ptcls
9.7 A 1 ess
15425 ptcls
9.5 A 1 ess
14943 ptcls
8.6 A 1 ess
14668 ptcls
9.7 A 2 ess
13231 ptcls
15.3 A 2 ess
13148 ptcls
16.8 A 2 ess
12478 ptcls
6.9 A 2 ess
12454 ptcls
10.1 A 2 ess
12452 ptcls
11.3 A 2 ess
12411 ptcls
11.1 A 2 ess
12204 ptcls
15.2 A 2 ess
12111 ptcls
12.3 A 1 ess
11949 ptcls
10.5 A 2 ess
11458 ptcls
15.7 A 2 ess
11372 ptcls
15.5 A 2 ess
11356 ptcls
12.7 A 2 ess
11203 ptcls
15.4 A 2 ess
10903 ptcls
11.8 A 2 ess
10485 ptcls
9.7 A 2 ess
10266 ptcls
17.6 A 2 ess
10202 ptcls
13.0 A 2 ess
10145 ptcls
14.1 A 3 ess
10144 ptcls
11.9 A 2 ess
10115 ptcls
11.5 A 2 ess
10108 ptcls
16.4 A 2 ess
10101 ptcls
12.3 A 2 ess
10053 ptcls
11.4 A 2 ess
10026 ptcls
12.8 A 2 ess
9818 ptcls
10.5 A 3 ess
9787 ptcls
16.5 A 2 ess
9729 ptcls
16.5 A 2 ess
9607 ptcls
14.1 A 2 ess
9565 ptcls
8.9 A 2 ess
9301 ptcls
12.0 A 2 ess
9163 ptcls
10.3 A 2 ess
9138 ptcls
16.3 A 3 ess
9030 ptcls
15.7 A 2 ess
8769 ptcls
12.2 A 2 ess
8747 ptcls
11.4 A 2 ess
8719 ptcls
14.7 A 3 ess
8644 ptcls
16.1 A 2 ess
8587 ptcls
10.7 A 2 ess
8539 ptcls
20.2 A 2 ess
8372 ptcls
16.3 A 3 ess
8348 ptcls
11.1 A 2 ess
8262 ptcls
10.0 A 3 ess
8257 ptcls
16.9 A 3 ess
8193 ptcls
15.0 A 2 ess
8097 ptcls
18.5 A 3 ess
8075 ptcls
17.4 A 2 ess
8050 ptcls
9.1 A 2 ess
8015 ptcls
13.2 A 2 ess
7957 ptcls
8.0 A 2 ess
7925 ptcls
15.0 A 2 ess
7860 ptcls
17.7 A 2 ess
7756 ptcls
17.0 A 2 ess
7709 ptcls
14.5 A 2 ess
7635 ptcls
10.5 A 2 ess
7553 ptcls
14.5 A 2 ess
7552 ptcls
15.8 A 2 ess
7523 ptcls
10.6 A 2 ess
7518 ptcls
8.8 A 2 ess
7481 ptcls
13.4 A 2 ess
7428 ptcls
16.4 A 2 ess
7427 ptcls
12.8 A 2 ess
7197 ptcls
16.0 A 2 ess
7083 ptcls
14.8 A 2 ess
6621 ptcls
13.1 A 3 ess
6108 ptcls
16.3 A 3 ess
5954 ptcls
18.8 A 2 ess
5804 ptcls
14.2 A 2 ess
5731 ptcls
11.1 A 2 ess
5728 ptcls
10.8 A 2 ess
5381 ptcls
16.0 A 2 ess

**Fig. S2**

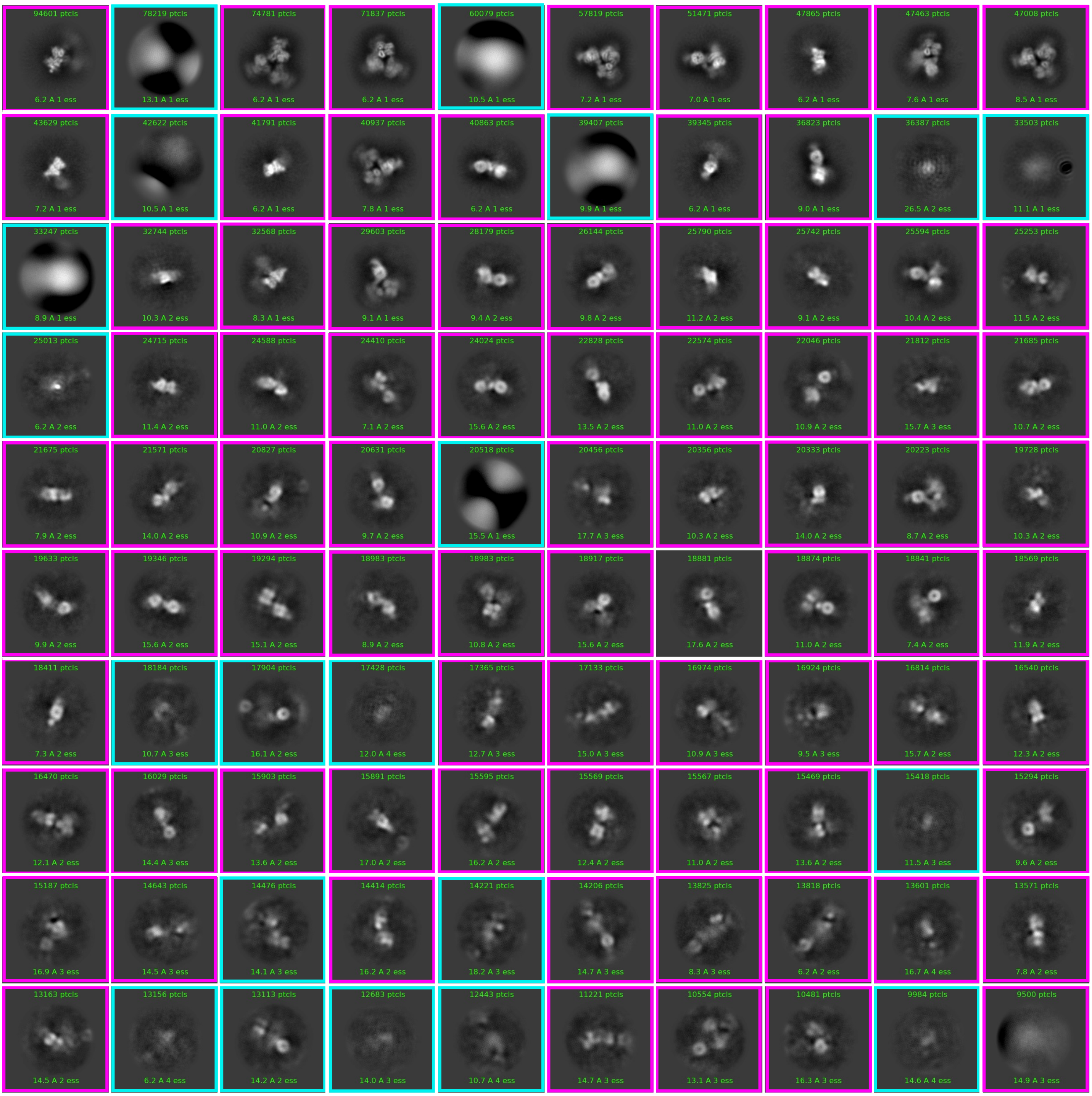
94601 ptcls
6.2 A 1 ess
78219 ptcls
13.1 A 1 ess
74781 ptcls
6.2 A 1 ess
71837 ptcls
6.2 A 1 ess
60079 ptcls
10.5 A 1 ess
57819 ptcls
7.2 A 1 ess
51471 ptcls
7.0 A 1 ess
47865 ptcls
6.2 A 1 ess
47463 ptcls
7.6 A 1 ess
47008 ptcls
8.5 A 1 ess
43629 ptcls
7.2 A 1 ess
42622 ptcls
10.5 A 1 ess
41791 ptcls
6.2 A 1 ess
40937 ptcls
7.8 A 1 ess
40863 ptcls
6.2 A 1 ess
39407 ptcls
9.9 A 1 ess
39345 ptcls
6.2 A 1 ess
36823 ptcls
9.0 A 1 ess
36387 ptcls
26.5 A 2 ess
33503 ptcls
11.1 A 1 ess
33247 ptcls
8.9 A 1 ess
32744 ptcls
10.3 A 2 ess
32568 ptcls
8.3 A 1 ess
29603 ptcls
9.1 A 1 ess
28179 ptcls
9.4 A 2 ess
26144 ptcls
9.8 A 2 ess
25790 ptcls
11.2 A 2 ess
25742 ptcls
9.1 A 2 ess
25594 ptcls
10.4 A 2 ess
25253 ptcls
11.5 A 2 ess
25013 ptcls
6.2 A 2 ess
24715 ptcls
11.4 A 2 ess
24588 ptcls
11.0 A 2 ess
24410 ptcls
7.1 A 2 ess
24024 ptcls
15.6 A 2 ess
22828 ptcls
13.5 A 2 ess
22574 ptcls
11.0 A 2 ess
22046 ptcls
10.9 A 2 ess
21812 ptcls
15.7 A 3 ess
21685 ptcls
10.7 A 2 ess
21675 ptcls
7.9 A 2 ess
21571 ptcls
14.0 A 2 ess
20827 ptcls
10.9 A 2 ess
20631 ptcls
9.7 A 2 ess
20518 ptcls
15.5 A 1 ess
20456 ptcls
17.7 A 3 ess
20356 ptcls
10.3 A 2 ess
20333 ptcls
14.0 A 2 ess
20223 ptcls
8.7 A 2 ess
19728 ptcls
10.3 A 2 ess
19633 ptcls
9.9 A 2 ess
19346 ptcls
15.6 A 2 ess
19294 ptcls
15.1 A 2 ess
18983 ptcls
8.9 A 2 ess
18983 ptcls
10.8 A 2 ess
18917 ptcls
15.6 A 2 ess
18881 ptcls
17.6 A 2 ess
18874 ptcls
11.0 A 2 ess
18841 ptcls
7.4 A 2 ess
18569 ptcls
11.9 A 2 ess
18411 ptcls
7.3 A 2 ess
18184 ptcls
10.7 A 3 ess
17904 ptcls
16.1 A 2 ess
17428 ptcls
12.0 A 4 ess
17365 ptcls
12.7 A 3 ess
17133 ptcls
15.0 A 3 ess
16974 ptcls
10.9 A 3 ess
16924 ptcls
9.5 A 3 ess
16814 ptcls
15.7 A 2 ess
16540 ptcls
12.3 A 2 ess
16470 ptcls
12.1 A 2 ess
16029 ptcls
14.4 A 3 ess
15903 ptcls
13.6 A 2 ess
15891 ptcls
17.0 A 2 ess
15595 ptcls
16.2 A 2 ess
15569 ptcls
12.4 A 2 ess
15567 ptcls
11.0 A 2 ess
15469 ptcls
13.6 A 2 ess
15418 ptcls
11.5 A 3 ess
15294 ptcls
9.6 A 2 ess
15187 ptcls
16.9 A 3 ess
14643 ptcls
14.5 A 3 ess
14476 ptcls
14.1 A 3 ess
14414 ptcls
16.2 A 2 ess
14221 ptcls
18.2 A 3 ess
14206 ptcls
14.7 A 3 ess
13825 ptcls
8.3 A 3 ess
13818 ptcls
6.2 A 2 ess
13601 ptcls
16.7 A 4 ess
13571 ptcls
7.8 A 2 ess
13163 ptcls
14.5 A 2 ess
13156 ptcls
6.2 A 4 ess
13113 ptcls
14.2 A 2 ess
12683 ptcls
14.0 A 3 ess
12443 ptcls
10.7 A 4 ess
11221 ptcls
14.7 A 3 ess
10554 ptcls
13.1 A 3 ess
10481 ptcls
16.3 A 3 ess
9984 ptcls
14.6 A 4 ess
9500 ptcls
14.9 A 3 ess

**Fig. S3**

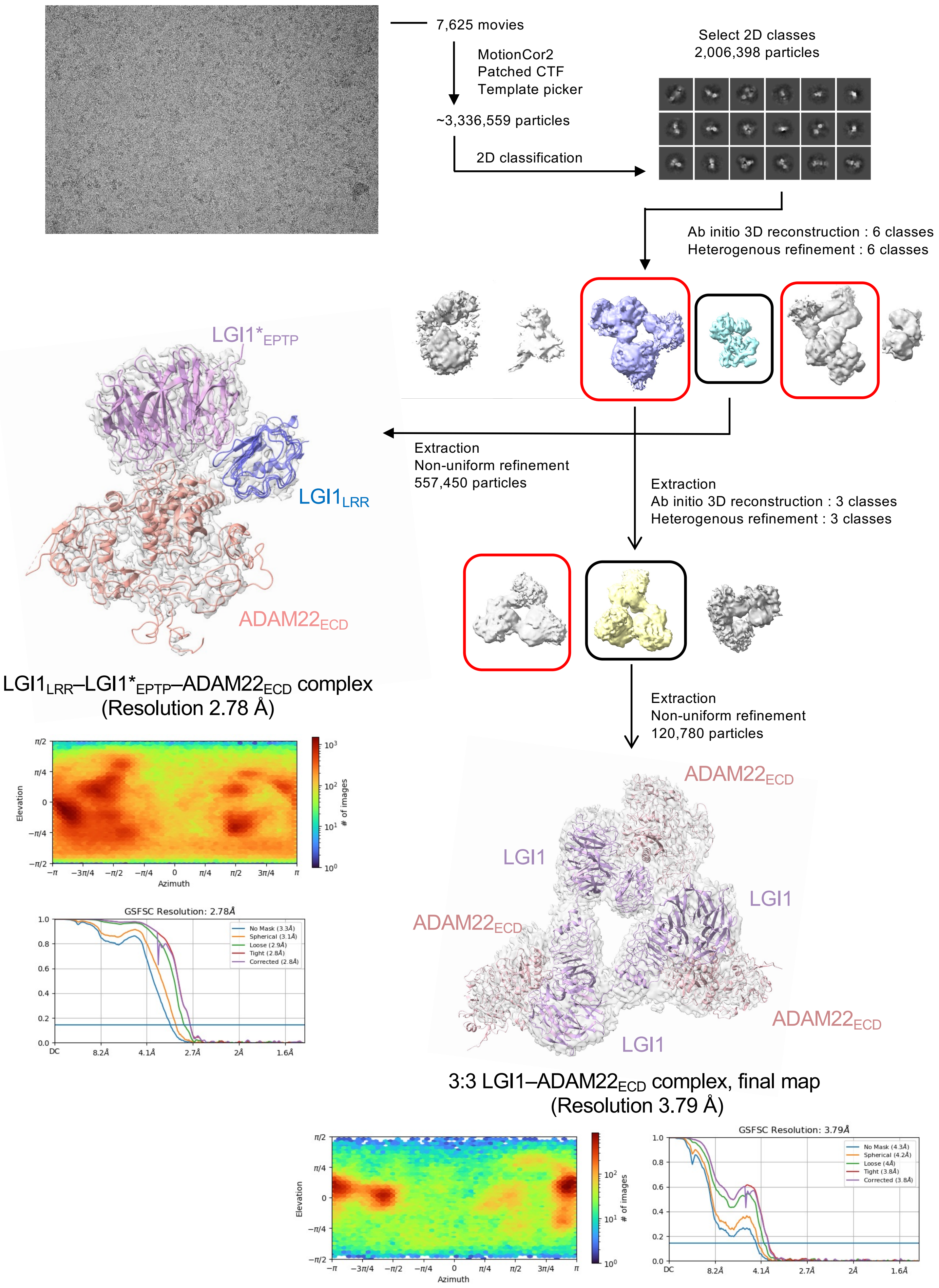

7,625 movies
MotionCor2
Patched CTF
Template picker
~3,336,559 particles
2D classification
Select 2D classes
2,006,398 particles
Ab initio 3D reconstruction : 6 classes
Heterogenous refinement : 6 classes
LGI1*EPTP
LGI1LRR
ADAM22ECD
Extraction
Non-uniform refinement
557,450 particles
Extraction
Ab initio 3D reconstruction : 3 classes
Heterogenous refinement : 3 classes
LGI1LRR–LGI1*EPTP–ADAM22ECD complex
(Resolution 2.78 Å)
Extraction
Non-uniform refinement
120,780 particles
Elevation
Azimuth
### of images
GSFSC Resolution: 2.78Å
No Mask (3.3Å)
Spherical (3.1Å)
Loose (2.9Å)
Tight (2.8Å)
Corrected (2.8Å)
ADAM22ECD
LGI1
LGI1
ADAM22ECD
ADAM22ECD
LGI1
3:3 LGI1–ADAM22ECD complex, final map
(Resolution 3.79 Å)
Elevation
Azimuth
### of images
GSFSC Resolution: 3.79Å
No Mask (4.3Å)
Spherical (4.2Å)
Loose (4Å)
Tight (3.8Å)
Corrected (3.8Å)

**Fig. S4**

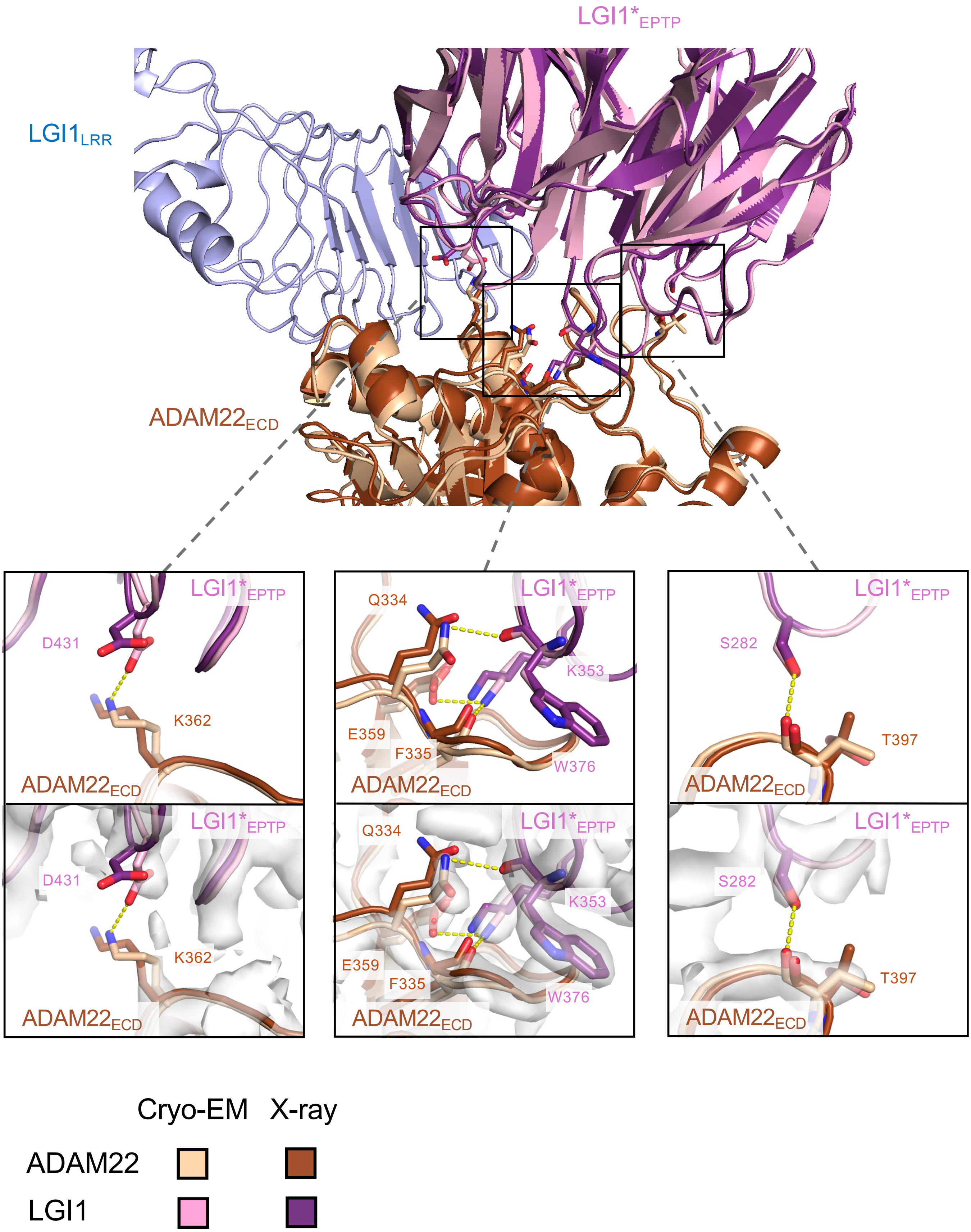

LGI1*EPTP
LGI1LRR
ADAM22ECD
LGI1*EPTP
D431
K362
ADAM22ECD
LGI1*EPTP
Q334
K353
E359
F335
W376
ADAM22ECD
LGI1*EPTP
S282
T397
ADAM22ECD
LGI1*EPTP
D431
K362
ADAM22ECD
LGI1*EPTP
Q334
K353
E359
F335
W376
ADAM22ECD
LGI1*EPTP
S282
T397
ADAM22ECD
Cryo-EM
X-ray
ADAM22
LGI1

**Fig. S5**

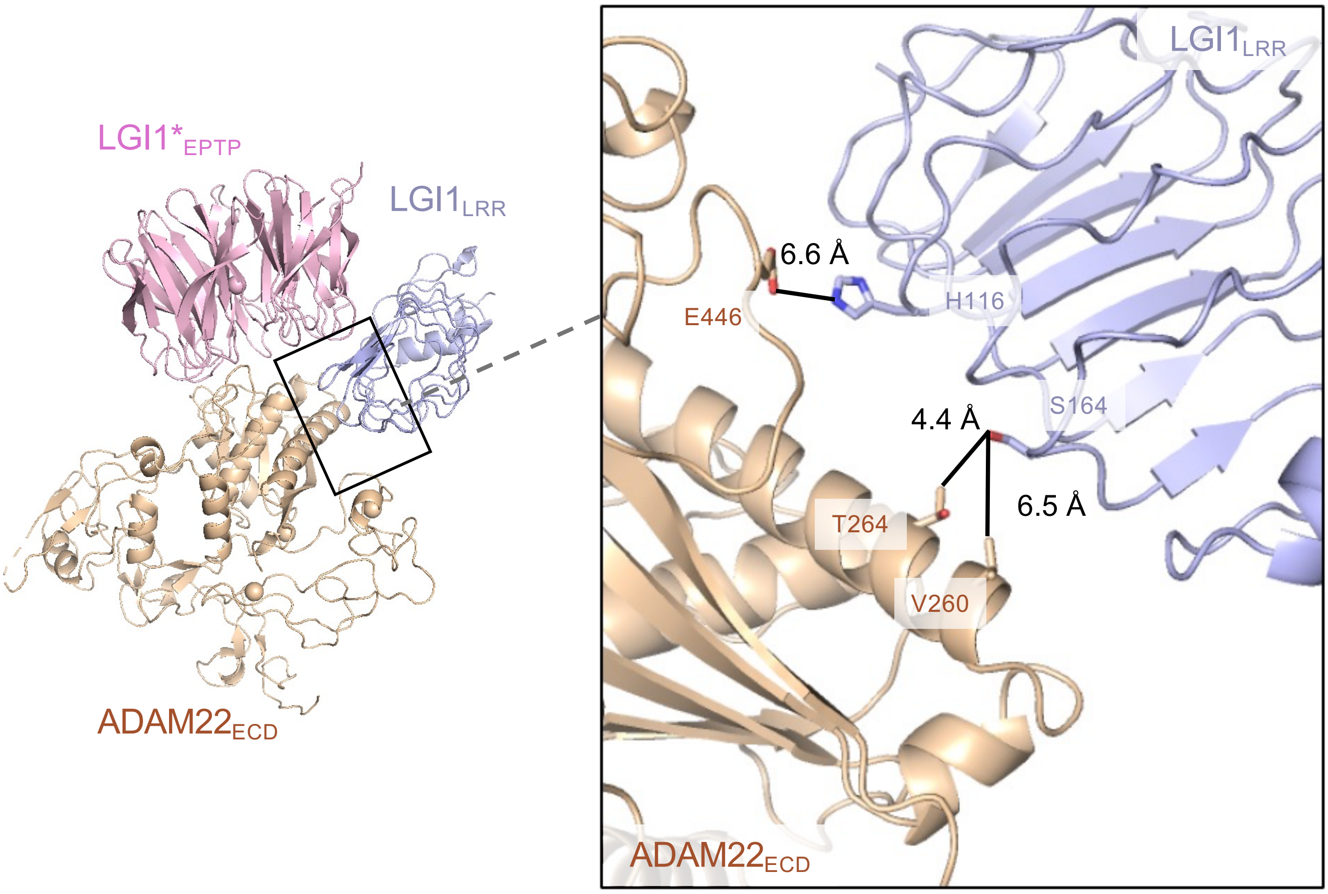

LGI1*EPTP
LGI1LRR
ADAM22ECD
LGI1LRR
6.6 Å
E446
H116
S164
4.4 Å
6.5 Å
T264
V260
ADAM22ECD

**Fig. S6**

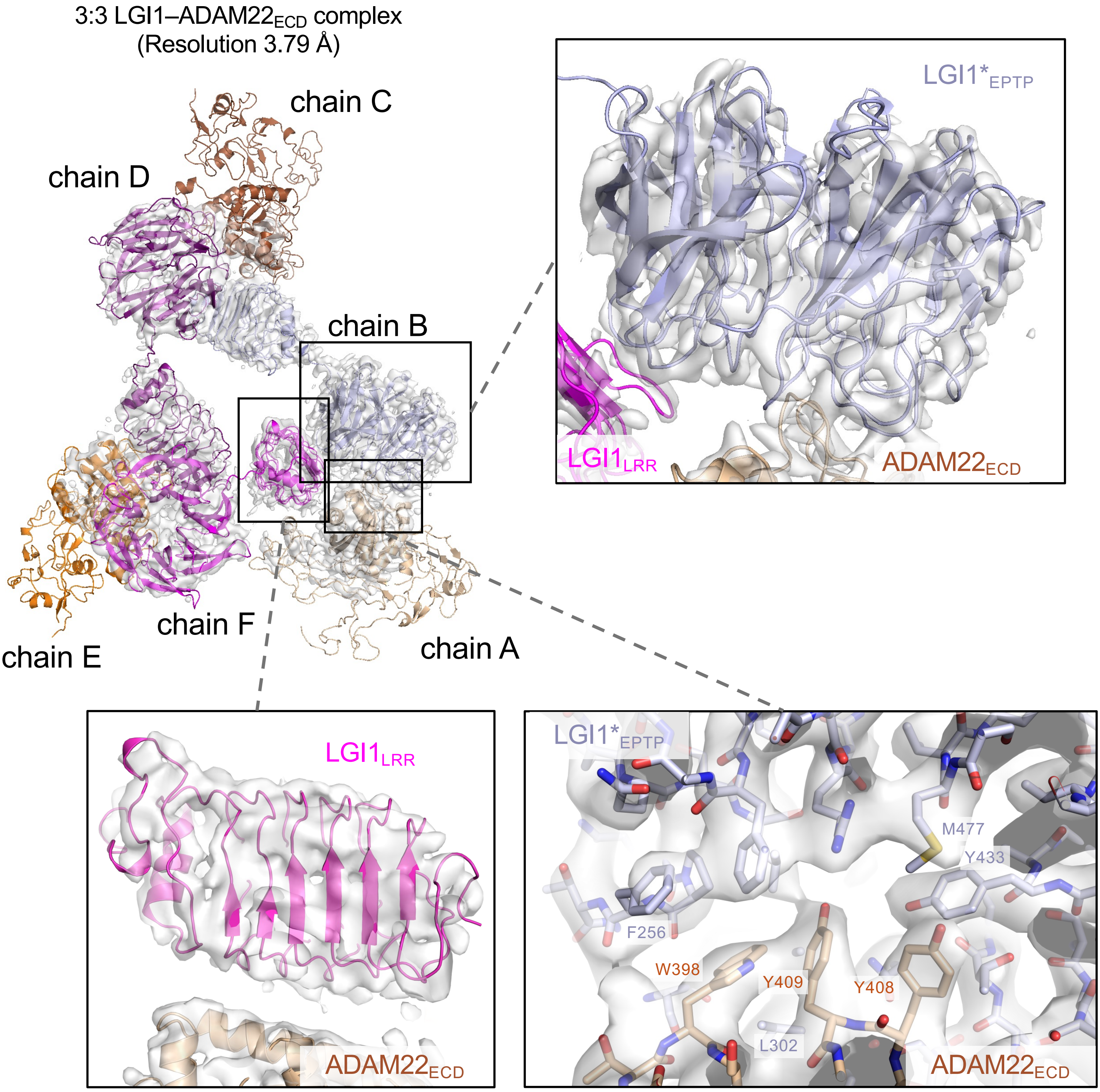

3:3 LGI1–ADAM22ECD complex
(Resolution 3.79 Å)
chain C
chain D
chain B
chain F
chain E
chain A
LGI1*EPTP
LGI1LRR
ADAM22ECD
LGI1LRR
ADAM22ECD
LGI1*EPTP
M477
Y433
F256
W398
Y409
Y408
L302
ADAM22ECD

**Fig. S7**

### 3:3 LGI1–ADAM22$_{ECD}$ complex

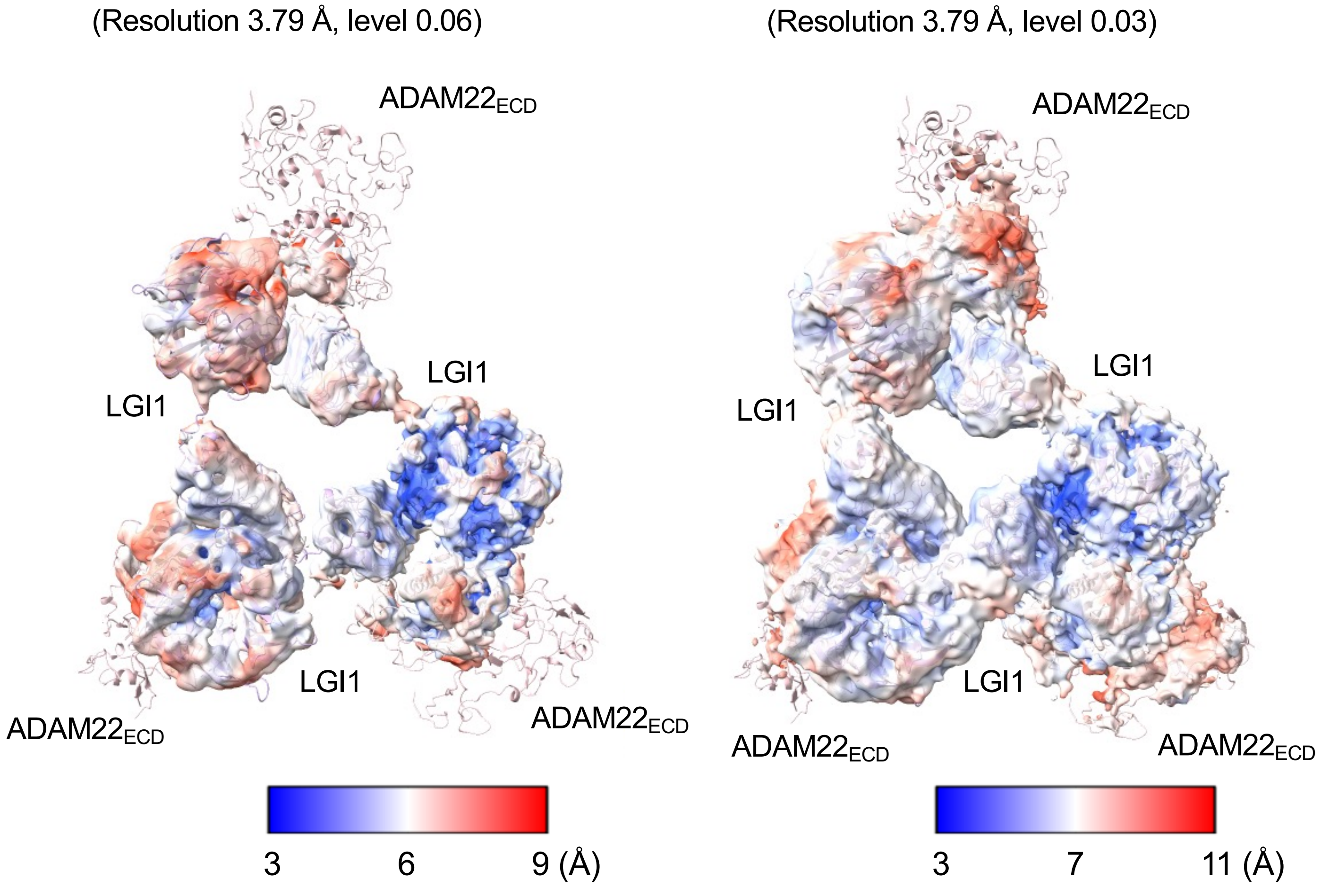

**Fig. S8**

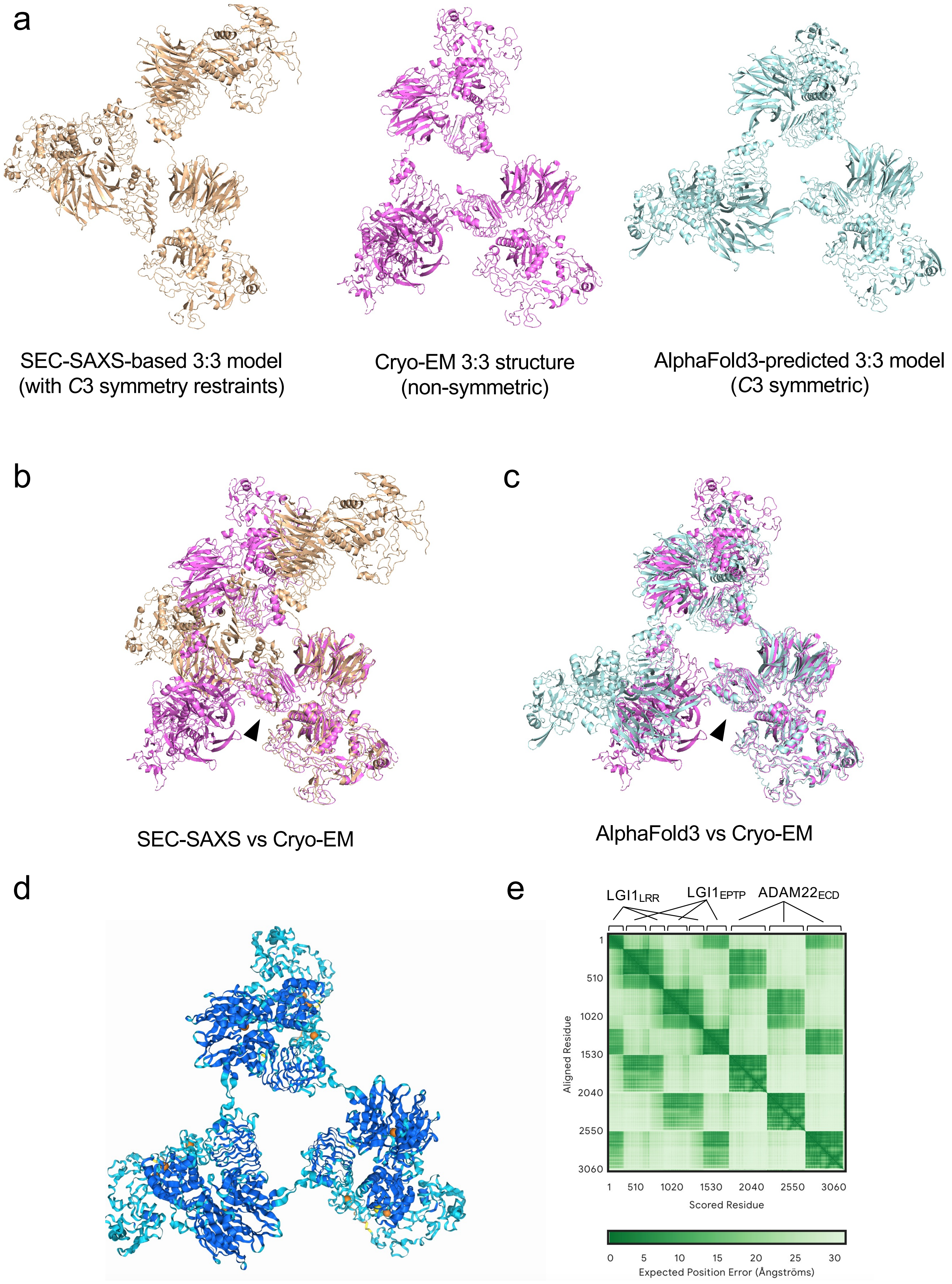
a
SEC-SAXS-based 3:3 model
(with C3 symmetry restraints)
Cryo-EM 3:3 structure
(non-symmetric)
AlphaFold3-predicted 3:3 model
(C3 symmetric)
b
SEC-SAXS vs Cryo-EM
c
AlphaFold3 vs Cryo-EM
d
e
LGI1LRR
LGI1EPTP
ADAM22ECD
Aligned Residue
1
510
1020
1530
2040
2550
3060
Scored Residue
Expected Position Error (Ångströms)
0
5
10
15
20
25
30

**Fig. S9**

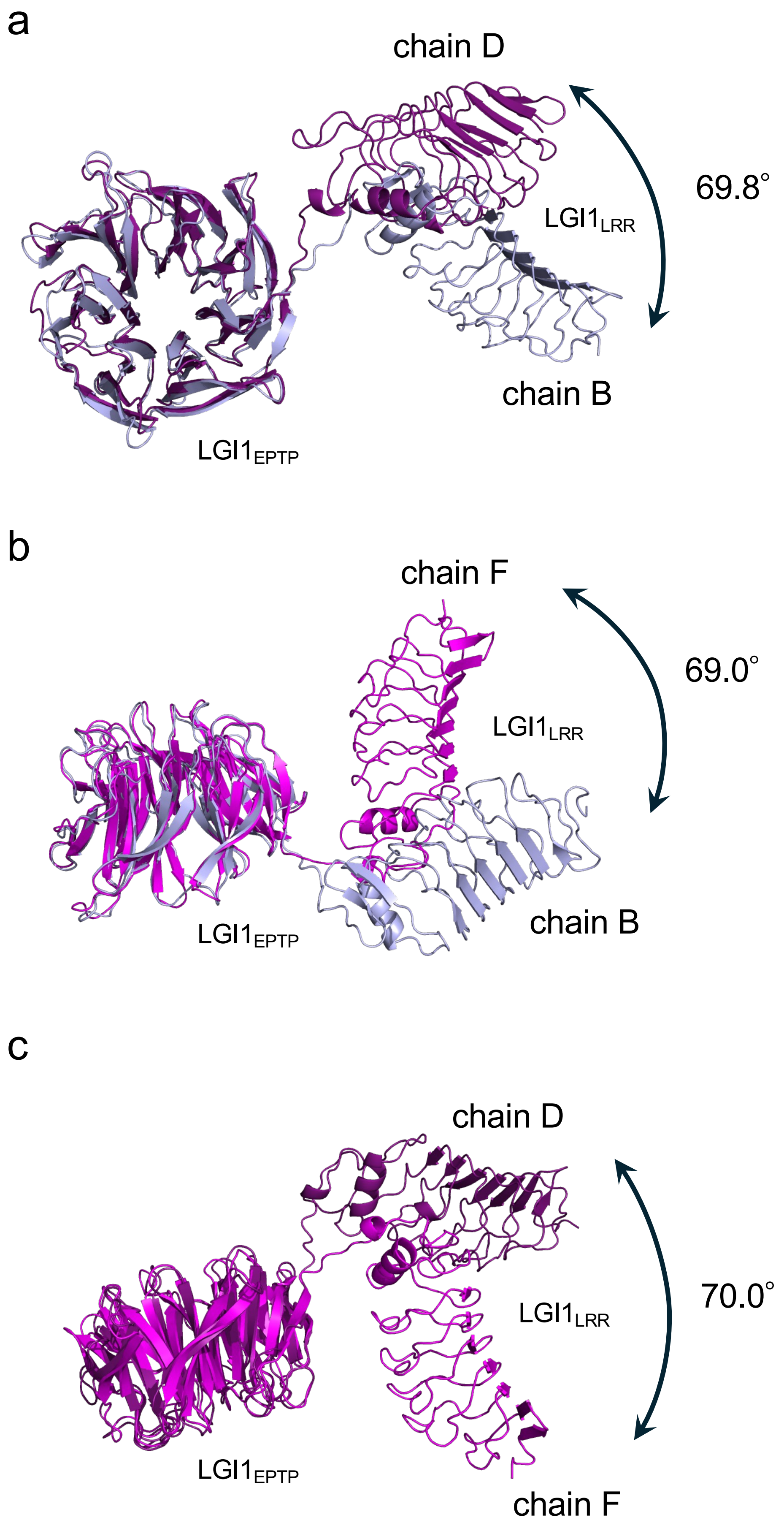
a
chain D
69.8°
LGI1LRR
chain B
LGI1EPTP
b
chain F
69.0°
LGI1LRR
LGI1EPTP
chain B
c
chain D
70.0°
LGI1LRR
LGI1EPTP
chain F

**Fig. S10**

**Data collection/processing and refinement statistics of cryo-EM single particle analysis**

| | $LGI1_{LRR}$–$LGI1^*_{EPTP}$–$ADAM22_{ECD}$ complex | 3:3 LGI1–$ADAM22_{ECD}$ complex |
|---|---|---|
| **Data collection and processing** | | |
| Magnification | 60,000 | 60,000 |
| Voltage (kV) | 300 | 300 |
| Dose rate ($e^-$/pixel/s) | 10.6743 | 10.6743 |
| Defocus range (μm) | –0.8 - –2.2 | –0.8 - –2.2 |
| Pixel size (Å) | 0.752 | 0.752 |
| Symmetry imposed | *C*1 | *C*1 |
| Initial particle images (no.) | 2,061,420 | 2,061,420 |
| Final particle images (no.) | 557,450 | 120,728 |
| Map resolution (Å) | 2.78 | 3.78 |
| FSC threshold | 0.143 | 0.143 |
| **Refinement** | | |
| Model resolution (Å) | | |
| FSC 0.143, unmasked/masked | 2.76/2.73 | 3.91/3.81 |
| Model composition | | |
| Non-hydrogen atoms | 7,846 | 23,526 |
| Protein residues | 993 | 2,979 |
| Ligands | 4 | 0 |
| *B* factors ($Å^2$) | | |
| Protein ($Å^2$) | 58.78 | 43.80 |
| Ligands ($Å^2$) | 51.14 | |
| RMS deviation | | |
| Bond length (Å) | 0.002 | 0.012 |
| Bond angle (°) | 0.474 | 1.153 |
| Molprobity score | 1.45 | 2.28 |
| Clash score | 6.47 | 24.79 |
| Rotamer outliers (%) | 0.00 | 0.60 |
| Ramachandran plot | | |
| Favored (%) | 97.56 | 94.16 |
| Allowed (%) | 2.44 | 5.84 |
| Disallowed (%) | 0.00 | 0.00 |

**Table S1**

**Domain motion analysis of LGI1 in the 3:3 LGI1–ADAM22$_{ECD}$ complex by the DynDom server.**

| Chain IDs | DynDom parameters | LGI1 |
|---|---|---|
| B vs D | Fixed domain | Residues 225-549 (RMSD 1.79 Å) |
| | Moving domain | Residues 43-224 (RMSD 1.16 Å) |
| | Rotation angle (°) | 69.8 |
| | Translation (Å) | –1.8 |
| | Closure (%) | 13.8 |
| | Bending residues | 215-225 |
| B vs F | Fixed domain | Residues 223-549 (RMSD 1.01 Å) |
| | Moving domain | Residues 43-222 (RMSD 1.19 Å) |
| | Rotation angle (°) | 69.0 |
| | Translation (Å) | -0.8 |
| | Closure (%) | 52.8 |
| | Bending residues | 215-223 |
| D vs F | Fixed domain | Residues 224-549 (RMSD 1.82 Å) |
| | Moving domain | Residues43-223 (RMSD 1.24 Å) |
| | Rotation angle (°) | 70.0 |
| | Translation (Å) | -1.7 |
| | Closure (%) | 99.0 |
| | Bending residues | 218-225 |

**Table S2**
